# In-silico immune cell deconvolution of the airway proteomes of infants with pneumonia reveals a link between reduced airway eosinophils and an increased risk of mortality

**DOI:** 10.1101/840090

**Authors:** Charles J Sande, Jacqueline M Waeni, James M Njunge, Martin N Mutunga, Elijah Gicheru, Nelson K Kibinge, Agnes Gwela

**Affiliations:** KEMRI-Wellcome Trust Research Programme, Kilifi, Kenya

**Keywords:** Respiratory syncytial virus, proteome, pneumonia, deconvolution

## Abstract

**Rationale:** Pneumonia is a leading cause of mortality in infants and young children. The mechanisms that lead to mortality in these children are poorly understood. Studies of the cellular immunology of the infant airway have traditionally been hindered by the limited sample volumes available from the young, frail children who are admitted to hospital with pneumonia. This is further compounded by the relatively low frequencies of certain immune cell phenotypes that are thought to be critical to the clinical outcome of pneumonia. To address this, we developed a novel in-silico deconvolution method for inferring the frequencies of immune cell phenotypes in the airway of children with different survival outcomes using proteomic data.

**Methods:** Using high-resolution mass spectrometry, we identified > 1,000 proteins expressed in the airways of children who were admitted to hospital with clinical pneumonia. 61 of these children were discharged from hospital and survived for more than 365 days after discharge, while 19 died during admission. We used machine learning by random forest to derive protein features that could be used to deconvolve immune cell phenotypes in paediatric airway samples. We applied these phenotype-specific signatures to identify airway-resident immune cell phenotypes that were differentially enriched by survival status and validated the findings using a large retrospective pneumonia cohort.

**Main Results:** We identified immune-cell phenotype classification features for 33 immune cell types. Eosinophil-associated features were significantly elevated in airway samples obtained from pneumonia survivors and were downregulated in children who subsequently died. To confirm these results, we analyzed clinical parameters from >10,000 children who had been admitted with pneumonia in the previous 10 years. The results of this retrospective analysis mirrored airway deconvolution data and showed that survivors had significantly elevated eosinophils at admission compared to fatal pneumonia.

**Conclusions:** Using a proteomics bioinformatics approach, we identify airway eosinophils as a critical factor for pneumonia survival in infants and young children.

## Introduction

Pneumonia is a leading cause of paediatric mortality word-wide. A recent study on the global burden of paediatric pneumonia conducted in seven countries found that viruses account for about 61% of all paediatric pneumonia infections, while about 27% of infection were attributed to bacterial pathogens, with RSV accounting for the largest etiological fraction of paediatric pneumonia^1^. More than 90% of the deaths that occur due to pneumonia in children under 5, occur in low resource settings, mainly due to the lack of paediatric intensive care facilities^2^. Very young infants especially those with comorbidities such as HIV and malnutrition have a poor survival prognosis following pneumonia infection. HIV-infected infants who develop a pneumonia infection are up to 10 times more likely to die from the infection than non-HIV infected children^3^, while those with malnutrition are more than 15 times more likely to die after admission^4^. The damage to the lungs caused by severe pneumonia appears to persist even after discharge from hospital, with recent estimates showing that post-discharge mortality in African children previously admitted to hospital with pneumonia being eight times greater than those discharged with other diagnoses^5^.

The difference in the immunological response to pneumonia between children who succumb to infection and those who survive it is not clear. An improved understanding of the immunobiology of this elevated post infection mortality risk will be crucial in identifying prognostic biomarkers of poor outcome, which will be critical in guiding care decisions in the first critical hours after admission. Most studies on the mechanisms of severe pneumonia in infants have been done using blood samples^6^, and whist these studies have provided significant insights into disease pathology, they might not fully recapitulate the immune response to infection in the airway. In the case of RSV, there has only been one study in the last 60 years that examined the lung samples of children who died from pneumonia. Archived post-mortem lung tissue from three children who died in the pre-intensivist era in the USA in the 1930s and 1940s was evaluated and airway obstruction with necrotic cells, fibrin, mucus, and fluid identified as a prominent feature in RSV lung infection^7^. Staining for immune cell populations was not technically feasible in these long term archival samples.

Due to the paucity of mechanistic data on the immunological response to pneumonia in the airway there is broad interest in understanding the dynamics of airway-resident immune cells following pneumonia infection in infants. This effort has been hindered by the unsuitability of routine airway sampling techniques for conventional cytometric analyses. Samples from nasopharyngeal washings and naso- and oropharyngeal swabbing are the most common methods of sampling the airways of sick children, and whilst ideal for molecular diagnostics, they are less suitable for phenotyping of airway resident immune cells using conventional cytometry techniques. Sampling of the airway by these methods typically results in limited cell yields and the cells that are recovered are generally highly enriched for granulocytes, complicating detailed characterisation of less abundant phenotypes using traditional flow cytometry-based tools^8^. Recent analysis of the cellular composition of upper airway by flow cytometry showed that the typical abundances of critical effector cells like T Cells, B Cells, Mast cells, Dendritic cells and NK cells to be less than 0.5% of all airway cells, while granulocytes were present at a median frequency of >90%^9^.

A potential way to address these shortcomings is to monitor changes in marker proteins that are uniquely or predominantly expressed by specific immune cell phenotypes and to use this information to infer the dynamics of the underlying cell types. We have recently described a high-resolution mass-spectrometry-based proteomics approach for characterizing the total proteome of the infant airway to a depth of more than 1,000 proteins^10,11^. The airway proteome characterized by this technique represents an unbiased snapshot of the underlying cell populations and can be leveraged to infer changes in the frequencies of the contributing cell phenotypes. Here, we describe an in-silico immune cell phenotype deconvolution approach where we use protein markers that are uniquely or predominantly expressed by specific immune cell subsets to infer the dynamics of those phenotypes in airways of children who survived or died from pneumonia. Phenotype-classification features were derived from a previously published data set containing the individual proteomes of purified immune cell populations^12^ (deconvolution data set) and these were then applied to airway proteome data from children with different pneumonia outcomes (Figure 1 contains a graphical description of the study design). Using this information, we identified an eosinophil-related protein signature that was elevated in the airways of children who survived pneumonia but that was downregulated in fatal pneumonia and in well controls. We subsequently validated this association using a large retrospective pneumonia cohort of >10,000 children.

**Figure 1:**
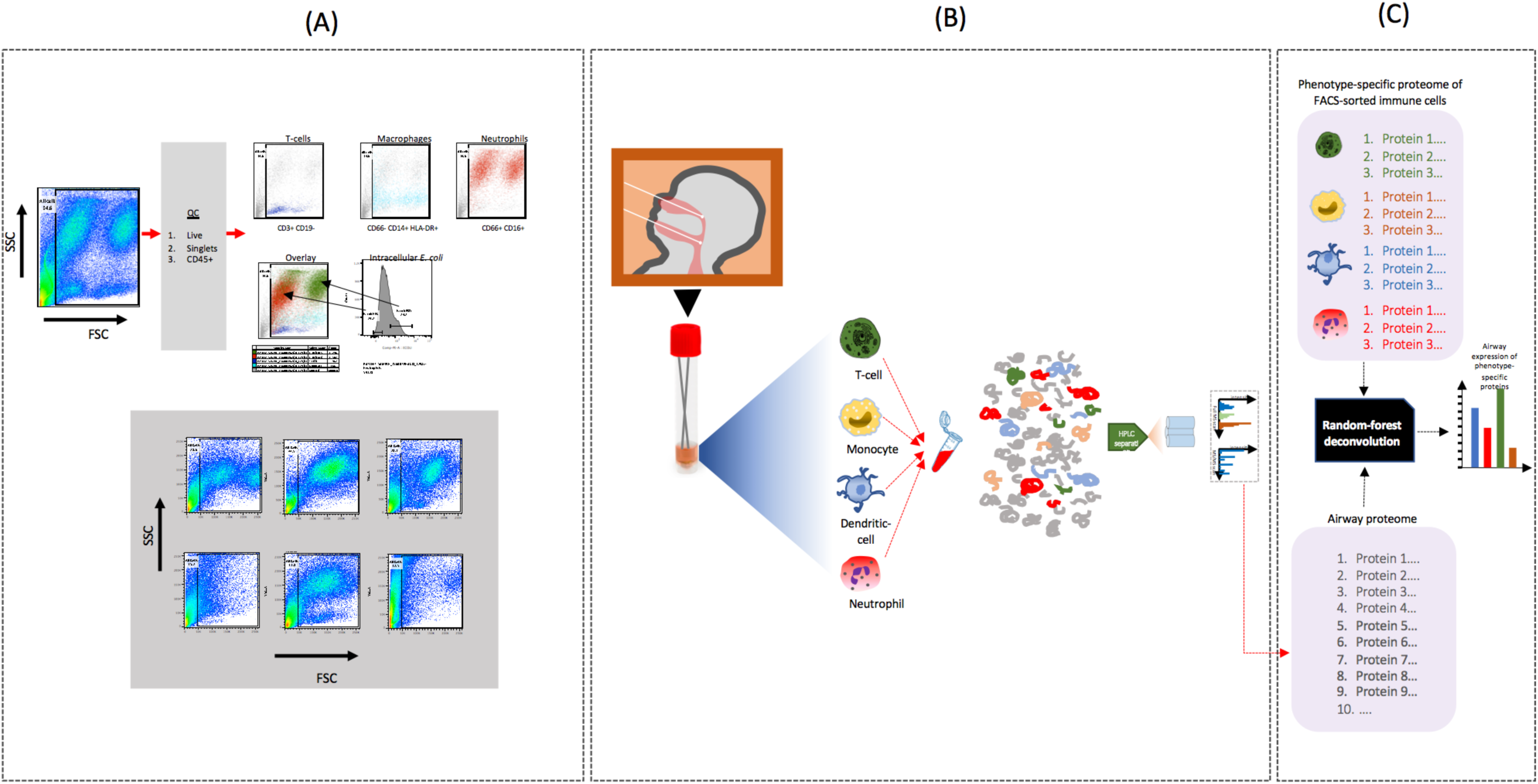
Analysis of the profile of airway-resident immune cells using flow cytometry and in-silico deconvolution. **(A)** To characterize the phenotype diversity of airway-resident immune cells, fresh naso- and oropharyngeal flocked swabs were obtained from children admitted to hospital with pneumonia and processed immediately by flow cytometry. Here the gating strategy is shown, where live, singlet cells were initially gated on CD45 expression, followed by fluorescent antibody staining for different cell surface markers. CD3+ and CD19-cells were gated as T-cells, CD66-, CD14+ and HLA-DR-cells were gated as macrophages/monocytes while neutrophils were gated on the basis of CD66 and CD16 double expression. The functional activity of neutrophils was characterized by co-culturing the cells with E.coli particles that were labelled with pHRhodo. Neutrophils that contained phagocytosed particles are shown in green in the overlay dot plot. The bottom represented FSC and SSC plots of samples obtained from 6 different infants and demonstrates a high level of diversity in the immune cell populations present in the airways of children with pneumonia. **(B)** In silico deconvolution analysis was based on high resolution mass spectrometry data obtained from the airway. Cells were obtained from the naso and oropharyngeal sites using separate swabs, which were both eluted in a common transport media. As shown in A above, these samples contained a broad diversity of immune cells including T-cells, monocytes/macrophages and neutrophils. These cells were isolated from the sample by centrifugation and processed for mass spectrometry analysis using a standard protocol (see methods). **(C)** In silico phenotype deconvolution was conducted initially by identifying phenotype specific protein features in a previously-published deconvolution data set using random forest feature selection. These features were then applied on airway proteome data set obtained from children admitted to hospital with severe pneumonia, and used to construct a detailed map of airway immunophenotypes and their associations with clinical outcomes.

## Results

To determine whether the deconvolution proteome data set contained sufficient resolution to distinguish individual immune cell phenotypes, we used protein expression levels from the data set to visualise phenotype segregation by nonmetric multidimensional scaling (NMDS). The analysis showed that major immune cell phenotypes including B-cells, T cells, natural killer (NK) cells, dendritic cells(DC), monocytes (MO), basophils, eosinophils and neutrophils could be distinctly segregated on the basis of differential protein expression (figure 2a). We then set out to identify individual proteins that could be used to accurately distinguish major and sub immune phenotypes (sub phenotypes defined as lower-level hierarchies of the major phenotypes – e.g. plasmacytoid and myeloid DCs within the major DC phenotype) within the deconvolution data set. Using the recursive feature elimination-paired random forest (RF) algorithm, boruta^13^, we identified classification features for 33 major and sub immune cell phenotypes (supplementary table 1). Since only a subset of these feature proteins are likely to be expressed within the specific context of the inflamed airway, we set out to determine which of the features identified by the model were also expressed in the airways of children admitted with pneumonia. For some cell types such as monocytes/macrophages and neutrophils, a substantial proportion (>30%) of the phenotype classification features from the feature selection model were also expressed in the airway, while for others (e.g. NK cells), a lower proportion of the classifiers were identified in the airway (figures 2B, D, F & H). Supplementary table 1 contains a complete list of all the identified features and their respective expression levels in the infant airway.

**Figure 2:**
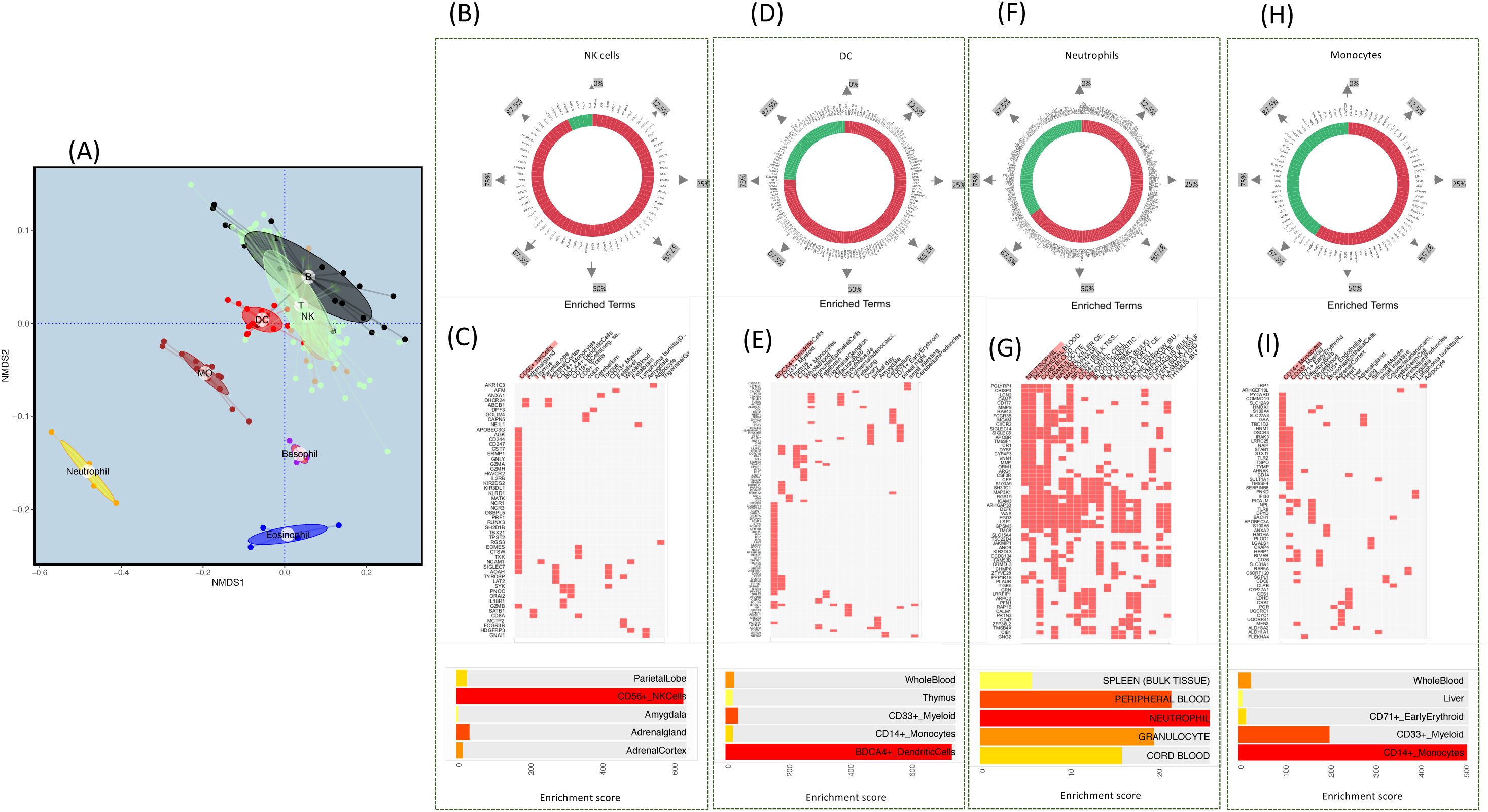
Use of proteomics data to segregate immune cell phenotypes. **(A)** A nonmetric multidimensional scaling (NMDS) plot was used to visualize the separation of immune cell phenotypes at the major phenotype level based on protein expression data from the deconvolution data set. We observed that expression profile of certain cell types including neutrophils, monocytes and basophils, resulted in a clear separation from other immune cell types. To identify specific proteins that could be used to distinguish phenotypes, we used random forest feature selection. Using this approach, we identified protein features for 33 phenotypes that could be used to disaggregate individual phenotypes from a complex mixture (Supplementary table 1). We then examined airway proteome data from children with pneumonia to determine whether any of the proteins identified were detectable in the airway. An example of the results is shown in **(B**,**D**,**F & H)**, where the phenotype classification features for NK cells, DCs, Neutrophils and Monocytes are shown labelled on the circular tile plot. The green tiles are the phenotype-specific feature proteins that were identified in the random forest model and that were also detected in the airway. We used enrichment analysis to determine whether the phenotype-classifier proteins identified by the random forest model for specific phenotypes could be used to recapitulate those the phenotypes in an unsupervised phenotype prediction analysis. The clustergrams shown in **(C**,**E**,**G & I)** depict a co-expression matrix of feature proteins and their known expression in different cell types. For 20 of the 33 phenotypes, (including the four examples shown here) the random forest phenotype classification, correctly predicted the original phenotype in the unsupervised prediction analysis. Each row on the clustergram represents a feature protein, while each colored box represents the known expression profile in a particular cell types. Horizonal panels at the bottom indicate enrichment scores from the unsupervised analysis.

Next, we undertook a more detailed analysis of the phenotype classification features that were identified by machine learning to determine whether they could be used to recapitulate the phenotypes in an unsupervised prediction analysis. We reasoned that the classification features of a particular phenotype would be broadly related to its functional properties and that when they are subjected to an independent unsupervised enrichment analysis, they would successfully predict the phenotypes from which they were initially derived. Using the unsupervised enrichment platform, enrichR^14,15^, we found that 20 of the 33 immune cell phenotypes were successfully predicted by their respective feature proteins (examples in figures 2 C, E, G & I). Subsequent analysis was restricted to these phenotypes. We visualised the performance of individual classification features by comparing their expression levels between different immune cell phenotypes. An example of this analysis is shown in figure 3A, where the monocyte classification feature, SERPINB2, is shown as being expressed at substantially higher levels in activated monocytes relative to all other cell phenotypes; the median expression level of SERPINB2 in classical activated monocytes was >10^4^ fold higher in monocytes, relative to all other phenotypes (figure 3B). Of the 64 classification features identified by machine learning for activated monocytes, 17 were detected in the airway samples from children with pneumonia (figure 3C).

**Figure 3:**
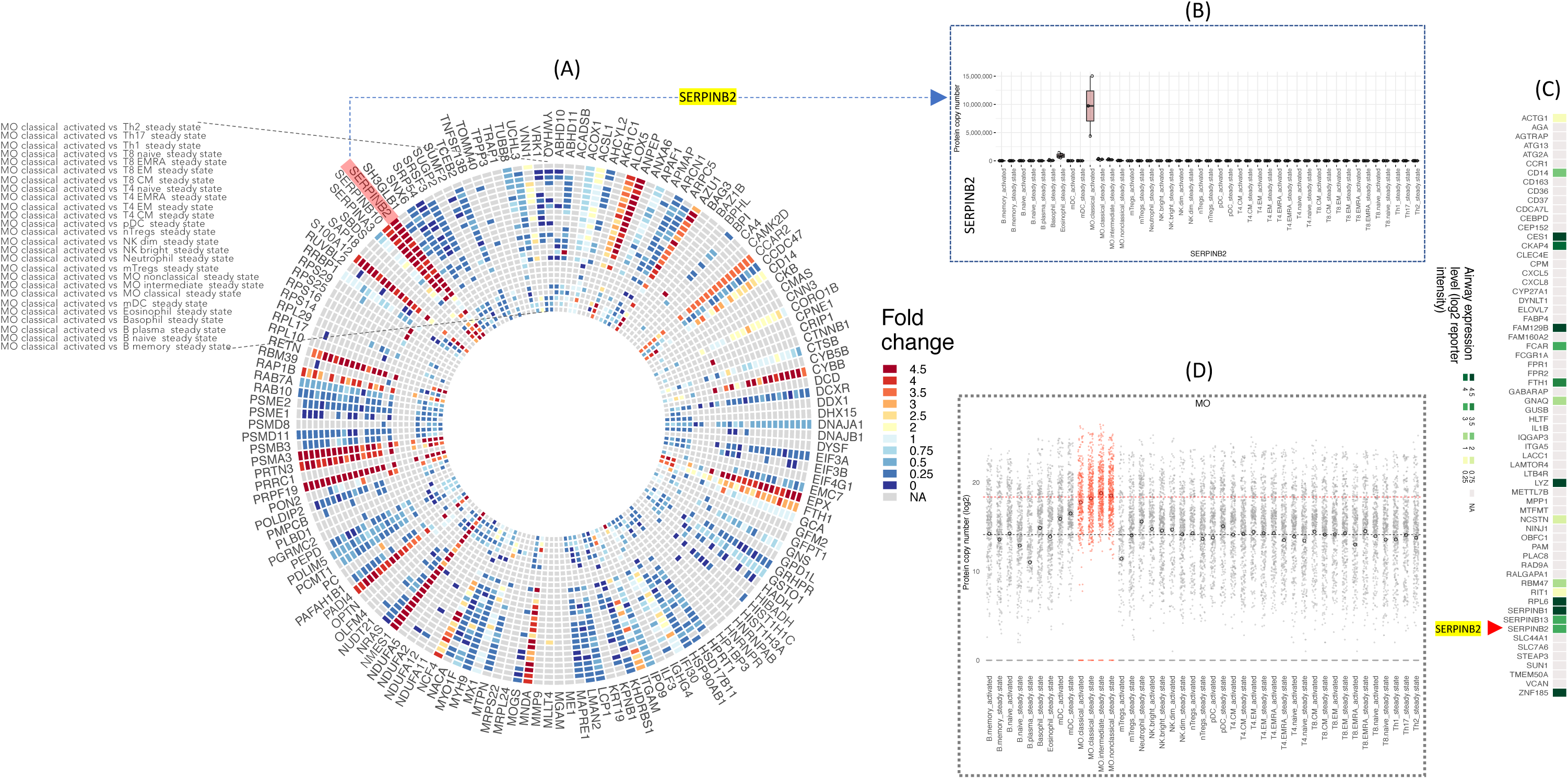
Detailed analysis of the phenotype-specific feature proteins. **(A)** We used a circular heat map to visualize differential expression of phenotype-specific proteins in a selection of immune cell phenotypes. In this example, the fold difference in protein expression between classical activated monocytes and other immune cell subsets is shown. Each spoke of the wheel represents a single protein, and each segment on a spoke represents the log_10_ fold difference between protein’s expression level in classical activated monocytes and each of the listed cell types. An increase in the intensity of the red hue indicates greater expression in classical activated monocytes, relative to the comparator phenotype. **(B)** Protein expression across different phenotypes: in this example, the expression level of a feature protein for activated monocytes (SERPINB2 – identified from the random forest model) was compared between different phenotypes and was seen to be expressed almost exclusively by monocytes. **(C)** The airway proteome data from paediatric pneumonia admissions was examined to determine whether any of the phenotype classification features for classical activated monocytes were also detected in the airways of children. Each segment represents a single feature, with proteins that were also detected in the airway depicted in a green hue. An increase in hue intensity, indicates increased airway expression for a particular protein. SERPINB2 (red arrow), was expressed at relatively high levels in the infant airway. **(D)** The expression level of all monocyte-specific features was plotted for monocytes and other cell types. Each black open circle denotes the expression level of a particular feature in other phenotypes, while red markers indicate their expression in monocytes. The dashed black and red lines indicated median expression of all features in monocytes and alternate cell types respectively.

To enhance the power of the identified features to resolve the constituent cell phenotypes of a complex sample, we generated a phenotype classification profile, in which all the classification features of a particular phenotype were aggregated and their combined expression was plotted relative to all other phenotypes. An example of this analysis is shown in figure 3D, where the combined expression of all monocyte-specific features (plotted in red) was compared to the expression level of the same proteins in all other phenotypes (plotted in gray). The results of this analysis showed that these proteins were expressed at a significantly higher level in monocytes compared to all other phenotypes (p<0.0001). Similar analysis for all other phenotypes is presented in supplementary figure 1. We then used t-SNE dimensional reduction analysis to visualise the segregation of major and sub phenotypes on the basis of the phenotype classification features (figure 4A). The results of this analysis showed that the feature classifier proteins could clearly distinguish most immune cell types; phenotypes like plasma cells (B.plasma), pDC, mDC, eosinophils, basophils, neutrophils and different monocyte sub phenotypes were clearly distinguishable from the rest of the phenotypes. Some sub-phenotypes such as central memory CD4 T cells (T4.CM) and effector memory CD4 T cells (T4.EM), could not be clearly disaggregated on the basis of feature expression alone.

**Figure 4:**
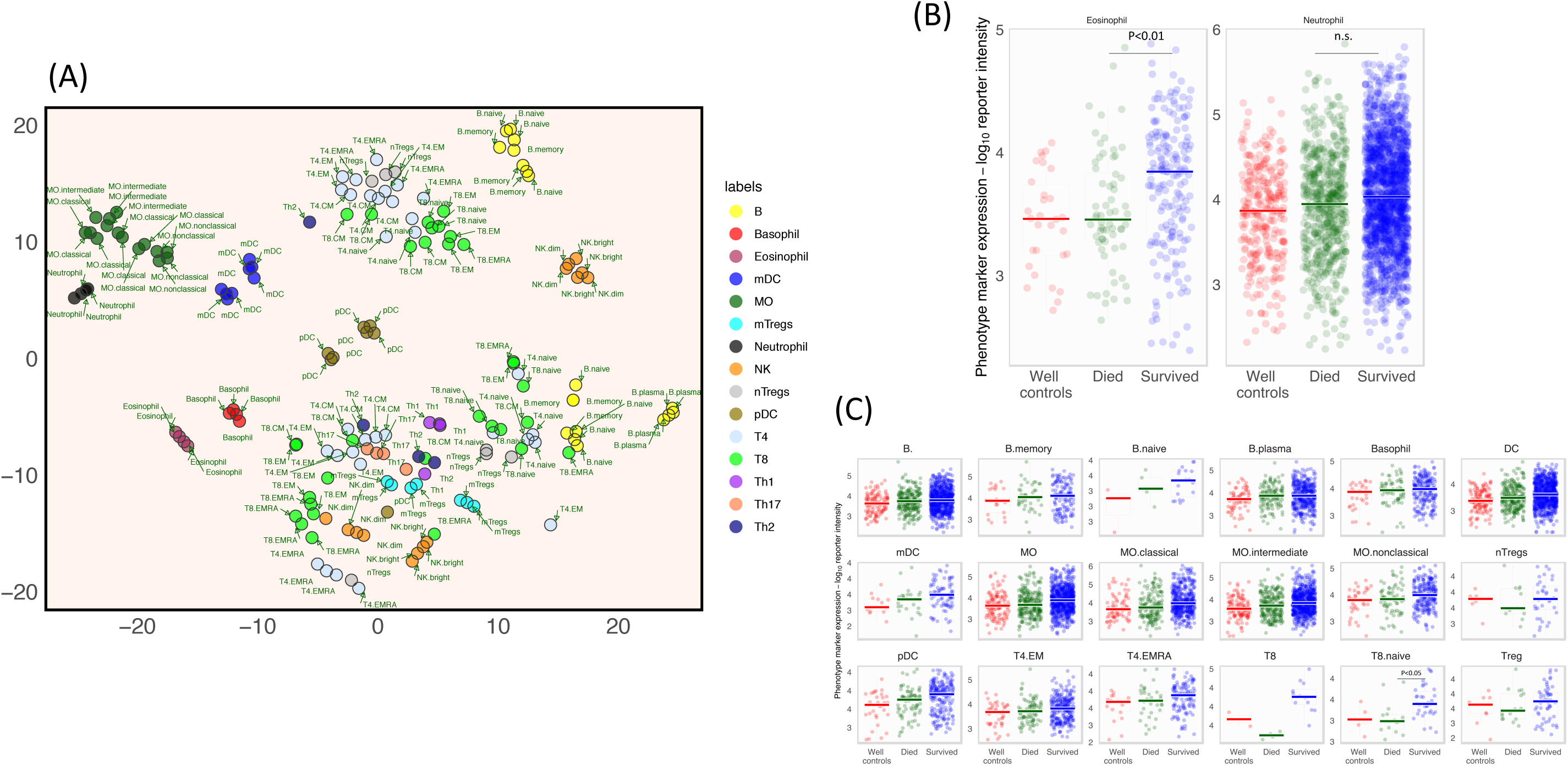
**(A)** Performance of all phenotype specific protein features in segregating different immune phenotypes and sub phenotypes was visualized using a t-SNE plot. Most phenotype classification features were able to disaggregate different phenotype hierarchies in the 2D t-SNE space. Neutrophils, eosinophils, monocytes, NK cells, dendritic cells, basophils among others were clearly separated from all other phenotypes. There was less clear segregation for T-cell sub phenotypes, including CD8, CD4 and regulatory T-cell phenotypes. **(B)** the expression of eosinophil- and neutrophil-specific features was compared between children with different survival outcomes of pneumonia as well as in well controls. Eosinophil features were expressed at significantly higher levels in survivors compared with non-survivors and well controls. In contrast, there was no difference in the expression of neutrophil features between these groups. **(C)** Feature classifiers for other immune cell phenotypes were also compared by survival status. No significant difference in expression by survival status was observed, with the exception of the CD8 (T8) phenotype, whose features were significantly upregulated in the airways of survivors relative to non-survivors and well controls.

We then applied the phenotype classification features identified by machine learning to airway proteome data obtained from naso/oropharyngeal samples from children who were admitted to hospital with clinical pneumonia and who either survived and were discharged from hospital(N=61) or ultimately died (N=19) in the course of admission. We also obtained similar data from age-matched well controls, who were sampled from the community (N=10). Using a false discovery rate (FDR) of 5%, we identified >1,000 proteins in the airways of these children. We then compared the expression levels of phenotype-specific feature proteins between these groups. We found that the expression levels of eosinophil-specific classification features were significantly overexpressed in children who survived infection, relative to non-survivors (P<0.0001) and controls (figure 4B). This was in contrast to neutrophils, where the expression level of phenotype-specific proteins did not vary by survival status (figure 4B). The expression level of proteins predominantly expressed by naïve CD8 T cells (T8.naive) were also significantly elevated in pneumonia survivors compared to non-survivors (P<0.05). We observed no significant differences for the remaining phenotypes (figure 4C). To validate these findings, we reviewed the clinical records of >10,000 children who had been admitted to Kilifi County Hospital Kenya over an approximate 10-year period with clinical pneumonia and for whom haematological data (including eosinophil and neutrophil counts) had been collected at admission. These data were stratified by survival status and the difference in eosinophils and neutrophil counts between these groups was tested. We found that eosinophil counts in children who survived pneumonia in the retrospective validation cohort (N=10,859) were significantly greater than those of non-survivors (N=1,604, p=0.0004) – Figure 5A, first panel. We then evaluated the risk of mortality in the first 10 days after admission in a subset of children who are at an elevated risk of pneumonia mortality (undernourished children), in order to determine the association between eosinophil counts and pneumonia survival in this group. Eosinophils counts in this group were stratified into two groups on the basis of median counts. Undernourished children who were admitted with pneumonia and with low (below median) eosinophil counts, were significantly more likely to die in the first 10 days of admission, relative to those with high eosinophil counts (Figure 5A, second panel). In contrast, no association with mortality was observed when neutrophils were analysed using a similar strategy (Figure 5B).

**Figure 5:**
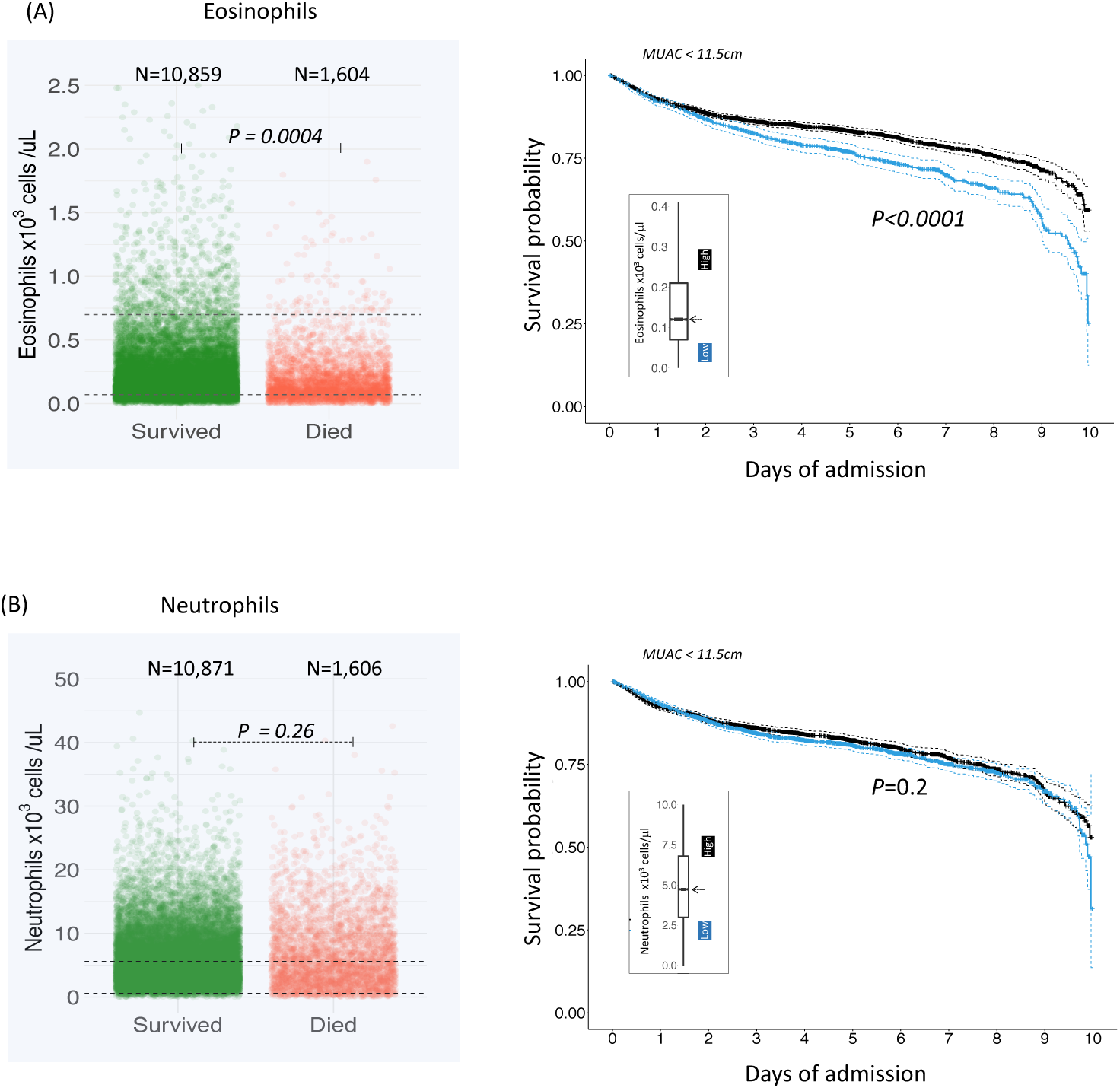
Validation of the phenotype associations with pneumonia mortality using a retrospective pneumonia cohort. **(A, first panel)** Eosinophils counts were analysed in a retrospective cohort of >10,000 children who had been admitted to hospital with clinical pneumonia and who either survived and were discharged or died during admission. Pneumonia survivors had significantly greater eosinophil counts at admission compared to non-survivors. **(A, second panel)** In a subset of children who were acutely undernourished at admission (defined as by a mid-upper arm (MUAC) circumference <11.5cm), children were stratified into two groups on the basis of the median eosinophil count. In the first 10 days after admission, children whose eosinophil counts were below the median were more likely to die relative to those with above-median counts. (B) In contrast, there was no association between neutrophil counts and survival status.

## Discussion

We report on a new method of deconvolving immune cell populations that are resident in the airways of infants and children with different survival outcomes of severe pneumonia. The study mucosal cellular immunity during very severe pneumonia has been hindered by a number of important hurdles including low sample volumes as well as the relatively low abundance of immune cell phenotypes that may be critical in directing the clinical course of pneumonia. In this study, we addressed these problems by using a machine learning approach to identify protein markers that could be used to deconvolve mixed immune cells. We then applied these markers to airway proteome data obtained from children with different survival outcomes of clinical pneumonia. Our results show that protein markers associated with eosinophils are elevated in the airways of survivors and are diminished in children who later died from infection. The airway levels of these eosinophil markers were no different between children who died and well controls, suggesting that the failure to mount an appropriate eosinophil response is a potential mechanism of pneumonia-related mortality, especially in high risk populations such as undernourished children. To validate the findings of the airway proteome analysis, we reviewed the hospitalisation records of >10,000 children who had been admitted to hospital in the previous 10 years with clinical pneumonia. The results of this retrospective analysis confirmed the observations made from analysis of the airway proteome and showed that children who died from pneumonia, had significantly lower blood eosinophil counts relative to those who survived. In contrast, neutrophil levels were not different between survivors and non survivors in both the airway proteome and in the retrospective validation cohort.

Previous studies have shown that eosinophils are activated in the airway shortly after pneumonia infection appear to contribute significantly to airway recovery. An increase in the expression of eosinophil-related markers in children with severe pneumonia has been associated with a reduced requirement for supplemental oxygen^16^, suggesting that these cells are a critical component in the host’s response to infecting pathogens in the airway. Airway-resident eosinophils have not been extensively characterised in previous pneumonia studies, although proteins expressed predominantly by eosinophils including leukotriene C4, eosinophil-derived neurotoxin (RNASE2) and eosinophil cationic protein (ECP) are expressed at high levels during viral pneumonia^17-19^. Taken together with the results of the current study, there appears to be a clear role for eosinophils in the response to pneumonia infection and that the failure to sufficiently recruit them to the airway may be a significant factor in mortality. A potential limitation of our study is that it was carried out using samples collected from the upper airway and not the lung. While lung samples would have undoubtedly been more informative of responses at the site of disease, the collection of these samples is a highly invasive process and exposes children to substantial additional risk without providing additional diagnostic value above nasopharyngeal or peripheral blood sampling. As a result, these samples are generally only available in settings with paediatric intensive care facilities, and even here, they are generally collected in children with atypically severe disease.

In addition to eosinophils, previous studies have shown infections that cause severe pneumonia such as RSV, trigger a strong cellular response, characterised by the influx of innate immune cells^20,21^. The initial response to infection is characterised by the airway recruitment of neutrophils, which express markers such as CD11b (ITGAM) and neutrophil granule proteins such as neutrophil elastase (ELANE)^22,23^. Other innate immune cells such as NK cells which expressed granzyme B are recruited in the lower airway and can be detected in in the lungs of mechanically-ventilated children with very severe pneumonia^24,25^. In addition to these cells, both myeloid and plasmacytoid dendritic cells are typically recruited into the airways of children with pneumonia in the early stages of infection^24,26^ and exhibit an activated proinflammatory phenotype^24^. Adaptive immune cells including CD4+ and CD8+ T cells are present in airway samples for children with viral pneumonia^20,27^. During infection, airway frequencies of granzyme B-secreting, activated CD8 T cells is greater in the airways of children with severe viral pneumonia^25^. In the present study, we found that protein markers of different phenotypes of CD8 T cells were significantly elevated in the airways of children who survived pneumonia relative to those who died or well controls. Although data was not available for validation of these phenotypes in the retrospective cohort, the data suggests a role for these cells in pneumonia survival. Previous studies in mechanically ventilated children showed that the frequency of lung-associated T cells increased as children recovered from infection, suggesting that these cells are an important component of effective local immunity against pneumonia and that deficits in the airways of children with pneumonia is a significant risk factor for mortality. In summary, the present data identifies a new approach for characterising the cellular immune response to pneumonia in the airway and identified critical immune populations that appear to be critical for survival. Future studies should aim to replicate these findings in samples collected from the lower respiratory tract.

## Methods

### Study site and population

This study recruited 80 infants and children who were admitted to Kilifi County Hospital with clinical pneumonia, defined using World Health Organisation syndromic criteria. Nasal samples were collected from each child for proteomic analysis. The microbial etiology of pneumonia was determined using both blood cultures and using a 15-target multiplex PCR panel for the detection of respiratory syncytial virus (RSV -A & B), rhinovirus, parainfluenza virus (1, 2, 3 & 4) adenovirus, influenza (A, B & C), coronavirus (OC43 & e229), human metapneumovirus and *Mycoplasma pneumoniae*. Children with clinical pneumonia signs and a positive diagnostic result from any of these tests were included in the analysis. Children were stratified into the survival group if they were alive for at least 365 days after discharge(n=61) while the mortality group comprised of children who died within 72 hours of admission (n=19). In addition to these groups, we recruited 10 age-matched well controls as a comparator group. Written informed consent was sought from the parents and legal guardians of all children who were sampled in this study. Ethical approval for the conduct of this study was granted by the Kenya Medical Research Institute’s Scientific and ethical research unit (SERU). All study procedures were conducted in accordance with Good Clinical Laboratory Practise (GCLP) standards.

### Analysis of airway proteomes using mass spectrometry

Naso- and oropharyngeal swab samples were centrifuged at 17,000xg for 10 mins at 4°C to obtain cell pellets which were washed once using PBS and lysed by bead-vortexing for 10 minutes in cell lysis buffer (RLT, Qiagen, Germany). Proteins (as well as DNA and RNA) were then extracted from the lysate using the AllPrep DNA/RNA/Protein Mini Kit (Qiagen, Germany) following manufacturers instructions. The concentration of total protein obtained was determined using the Bradford assay (Bio-Rad, USA). Thirty micrograms (30µg) of total protein from each sample was then reduced with 10mM tris(2-carboxyethyl)phosphine (TCEP, Sigma-Aldrich, USA) at 55°C for 1h and subsequently alkylated with 18mM IAA (Sigma-Aldrich, USA) for 30 minutes at room temperature, while keeping the reaction protected from light. Proteins were precipitated overnight at −20°C with six volumes of pre-chilled (−20°C) acetone (Sigma-Aldrich, USA). The samples were centrifuged at 8,000xg for 10 minutes at 4°C to obtain the protein pellets and supernatants were discarded. The protein pellet was resuspended in 100µl of 50mM Triethylammonium bicarbonate (TEAB, Sigma-Aldrich, USA). Trypsin (Sigma-Aldrich, USA) was added to the protein samples at a trypsin-protein sample ratio of 1:10 and protein digestion was allowed to proceed overnight at 37°C with shaking. The peptide samples were randomly assigned to 10 individual batches: each containing nine patient samples and one pooled control sample. The pooled control sample consisted of a pool of peptides from all patient samples. The peptide samples derived from individual patients were then individually labelled using the TMT10plex mass tag kit (Thermo scientific, USA) according to manufacturer’s instructions, with one isobaric tag being exclusively used to label the pooled control sample. The labelled peptides for each 10plex were subsequently combined to generate 10 individual pools. The labelled peptide pools were desalted using P10 C18 pipette ZipTips (Millipore, USA) according to the manufacturer’s instructions. Eluted peptides were dried in a Speedvac concentrator (Thermo Scientific, USA). Peptides (8 μl) were loaded using a Dionex Ultimate 3000 nano-flow ultra-high-pressure liquid chromatography system (Thermo Scientific, USA) on to a 75µm × 2 cm C18 trap column (Thermo Scientific, USA) and separated on a 75µm × 50 cm C18 reverse-phase analytical column (Thermo Scientific) at heated at 40°C. For LFQ protein quantification; elution was carried out with mobile phase B (80% acetonitrile with 0.1% formic acid) gradient (4 to 30%) over 310 min at a flow rate of 0.25 μl/min. Each LC run was finished by washout with 98% B for 10 min and re-equilibration in 2% B for 30 min. Five blanks of 40 min each were run on the column between each injection comprising of two wash cycles with 90% B and an equilibration phase of 15 min to avoid sample carryover. Peptides were measured using a Q Exactive Orbitrap mass spectrometer (Thermo Scientific, USA) coupled to the chromatography system via a nano-electrospray ion source (Thermo Scientific). On the Q Exactive, the ms^1 settings for peptides were: Resolution, 70000; AGC target, 3e6; maximum IT, 120 ms; scan range, 400-1800 m/z; while the ms^2 settings for fragmentation spectra of peptides were: Resolution, 17000 (35000 for labelled peptides); AGC target, 5e4; maximum IT, 120 ms; isolation window, 1.6 m/z. MS data were acquired by data dependent acquisition where the top 12 (15 for labelled peptides) most intense precursor ions in positive mode were selected for ms^2 Higher-energy C-trap dissociation fragmentation which were subsequently excluded for the next 45 s following fragmentation event. Charge exclusion was set to ignore peptide spectrum matches that were unassigned, singly charged, and those with ≥+8 charges. Raw mass spectrometer files were analysed by MaxQuant software version 1.6.0.1. by searching against the human Uniprot FASTA database (downloaded February 2014) using the Andromeda search engine..

Airway samples were centrifuged at 17,000xg for 10 mins at 4°C to obtain cell pellets which were washed once using PBS and lysed by bead-vortexing for 10 minutes in cell lysis buffer (RLT, Qiagen, Germany). Proteins (as well as DNA and RNA) were then extracted from the lysate using the AllPrep DNA/RNA/Protein Mini Kit (Qiagen, Germany) following manufacturers instructions. The concentration of total protein obtained was determined using the Bradford assay (Bio-Rad, USA). Thirty micrograms (30µg) of total protein from each sample was then reduced with 10mM tris(2-carboxyethyl)phosphine (TCEP, Sigma-Aldrich, USA) at 55°C for 1h and subsequently alkylated with 18mM IAA (Sigma-Aldrich, USA) for 30 minutes at room temperature, while keeping the reaction protected from light. Proteins were precipitated overnight at −20°C with six volumes of pre-chilled (−20°C) acetone (Sigma-Aldrich, USA). The samples were centrifuged at 8,000xg for 10 minutes at 4°C to obtain the protein pellets and supernatants were discarded. The protein pellet was resuspended in 100µl of 50mM Triethylammonium bicarbonate (TEAB, Sigma-Aldrich, USA). Trypsin (Sigma-Aldrich, USA) was added to the protein samples at a trypsin-protein sample ratio of 1:10 and protein digestion was allowed to proceed overnight at 37°C with shaking. The peptide samples were randomly assigned to 10 individual batches: each containing nine patient samples and one pooled control sample. The pooled control sample consisted of a pool of peptides from all patient samples. The peptide samples derived from individual patients were then individually labelled using the TMT10plex mass tag kit (Thermo scientific, USA) according to manufacturer’s instructions, with one isobaric tag being exclusively used to label the pooled control sample. The labelled peptides for each 10plex were subsequently combined to generate 10 individual pools. The labelled peptide pools were desalted using P10 C18 pipette ZipTips (Millipore, USA) according to the manufacturer’s instructions. Eluted peptides were dried in a Speedvac concentrator (Thermo Scientific, USA). Peptides (8 μl) were loaded using a Dionex Ultimate 3000 nano-flow ultra-high-pressure liquid chromatography system (Thermo Scientific, USA) on to a 75µm × 2 cm C18 trap column (Thermo Scientific, USA) and separated on a 75µm × 50 cm C18 reverse-phase analytical column (Thermo Scientific) at heated at 40°C. For LFQ protein quantification; elution was carried out with mobile phase B (80% acetonitrile with 0.1% formic acid) gradient (4 to 30%) over 310 min at a flow rate of 0.25 μl/min. Each LC run was finished by washout with 98% B for 10 min and re-equilibration in 2% B for 30 min. Five blanks of 40 min each were run on the column between each injection comprising of two wash cycles with 90% B and an equilibration phase of 15 min to avoid sample carryover. Peptides were measured using a Q Exactive Orbitrap mass spectrometer (Thermo Scientific, USA) coupled to the chromatography system via a nano-electrospray ion source (Thermo Scientific). On the Q Exactive, the ms^1 settings for peptides were: Resolution, 70000; AGC target, 3e6; maximum IT, 120 ms; scan range, 400-1800 m/z; while the ms^2 settings for fragmentation spectra of peptides were: Resolution, 17000 (35000 for labelled peptides); AGC target, 5e4; maximum IT, 120 ms; isolation window, 1.6 m/z. MS data were acquired by data dependent acquisition where the top 12 (15 for labelled peptides) most intense precursor ions in positive mode were selected for ms^2 Higher-energy C-trap dissociation fragmentation which were subsequently excluded for the next 45 s following fragmentation event. Charge exclusion was set to ignore peptide spectrum matches that were unassigned, singly charged, and those with ≥+8 charges. Raw mass spectrometer files were analysed by <u>MaxQuant software</u> version 1.6.0.1. by searching against the human Uniprot FASTA database (downloaded February 2014) using the Andromeda search engine.

### Analysis of airway-resident immune cells using flow cytometry

1ml of nasopharyngeal and oropharyngeal swab samples obtained from children was centrifuged at 17,000xg for 7 minutes, after which 800⧅l of the supernatant was removed and discarded. The remaining 200μl were split into two aliquots of 100μl each. The first aliquot was used for neutrophil phenotyping assays and the other was used for neutrophil phagocytosis assays. 20μl of a pre-constituted cocktail of the following antibodies (from ThermoFisher) was used to label both aliquots – CD45, CD16, CD14, CD3, CD19, HLA-DR, CD66b, CD11b and a Live-dead marker. With the exception of the live/dead marker, all other antibodies were diluted 1:100 in FACS buffer. The live/dead marker was prepared at a 1:1000 dilution in FACS buffer. For the phagocytosis assay tube, 20⧅l of opsonised *Escherichia coli* (*E.coli*) was added to the tube (pHrodo Red E. coli BioParticles; ThermoFisher). The bacteria was initially prepared by mixing the *E.coli* strain with new-born calf sera followed by a 30-minute incubation at 37°C. After this step, both tubes were incubated at 37°C for 35 minutes. After the incubation, 20⧅l of a live-dead marker was added to each tube and incubated for 10 minutes at 37°C. The reaction was stopped by adding 500⧅l of 1X RBC lysis buffer followed by a 5-minute incubation. Cells in each tube were then spun down at 2,700xg for 1 minute and the supernatant discarded. Cells were then washed twice with FACS buffer, after which 350⧅l of FACS flow was added. Cells were then analysed immediately on a BD LSR Fortessa instrument. The following gating strategy was used to detect airway-resident neutrophils: Debris were excluded on the basis of their forward (FSC-A) and side (SSC-A) scatter characteristics, doublets were excluded using FSC-A versus FSC-H and dead cells were excluded using the live-dead marker. Data was analysed using FlowJo software.

### Statistical data analysis

Data were analyzed in R. deconvolution analysis by random forest classification was done using the Boruta package, with a maximum of 300 iterations. Protein classifiers that whose mean expression in the test phenotype was significantly greater than alternative phenotypes were taken forward for further analysis. T-SNE analysis was carried out using the Rtsne, with the iterations parameter set to a maximum of 300 and a perplexity value of 30. Analysis was carried out in two dimensional space. Cell type enrichment analysis was done one the enrichR platform. The input search term used for enrichment analysis was the RF protein classifier lists derived from random forest classification. All pairwise comparisons between the expression level of RF classifiers between phenotypes and frequency of immune cell types between survival states was done using t-tests on log10 normalised data. The deconvolution data set that was used to identify phenotype-specific protein classifiers was obtained from a previously published paper by Rieckmann et al.^12^

## Data availability

The proteomics data reported in this paper are available at the ProteomeXchange Consortium database (<u>Accession numbers:</u> PXD009403).

## Acknowledgements

This study was supported by fellowship funding to CJS from the Wellcome Trust (WT105882MA). The funder played no role in the conceptualization, design, data collection, analysis, decision to publish, or preparation of the manuscript

## Author contributions

CJS designed the study

CJS, JMN, MNM, ETG, AG conducted the experiments

CJS,NKK analysed the data

CJS wrote the manuscript

All authors reviewed and approved the manuscript

