## Supplementary figures and images for "In-silico immune cell deconvolution of the airway proteomes of infants with pneumonia reveals a link between reduced airway eosinophils and an increased risk of mortality"

### Supplementary figure

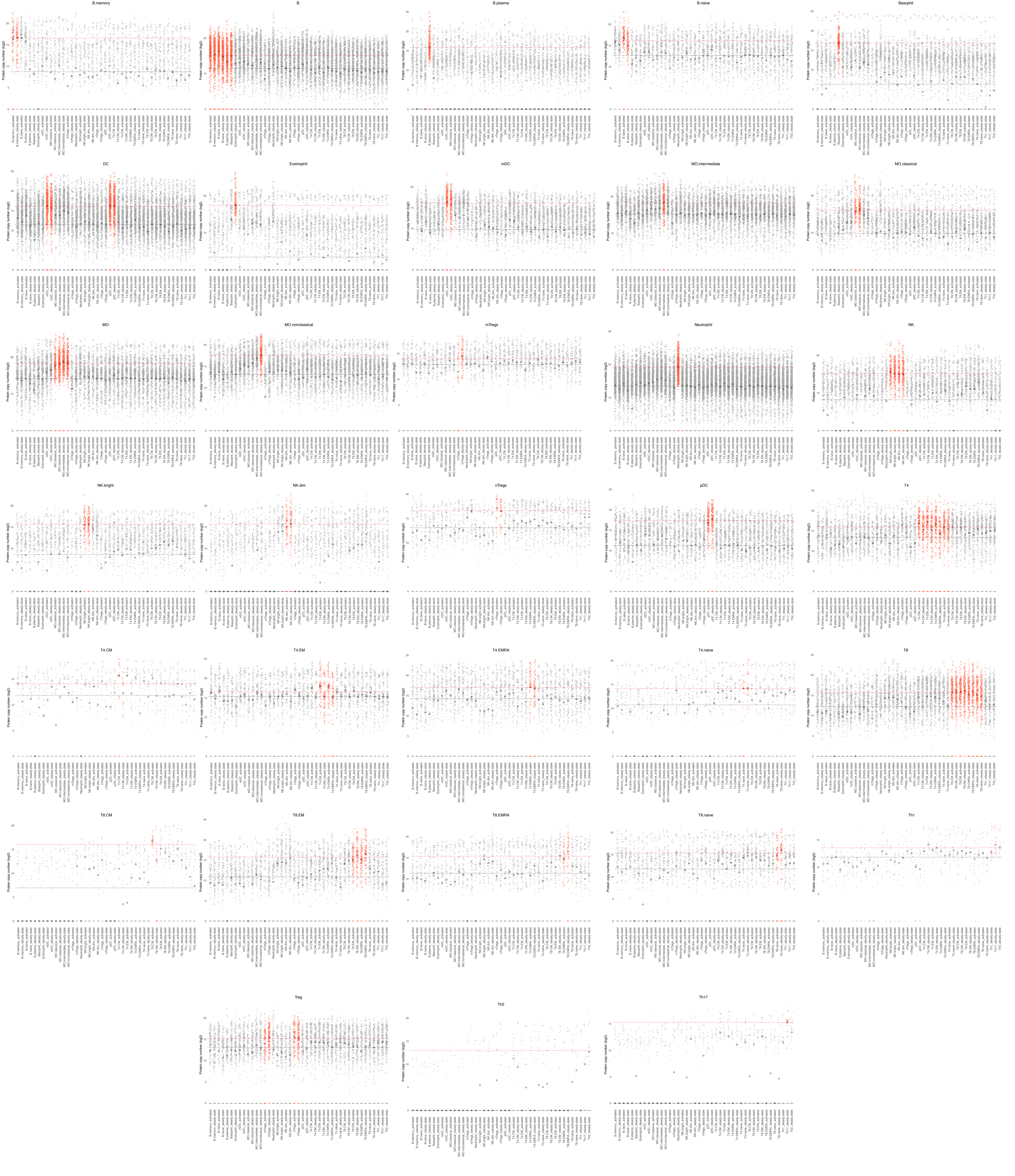
