## Supplementary table for "In-silico immune cell deconvolution of the airway proteomes of infants with pneumonia reveals a link between reduced airway eosinophils and an increased risk of mortality"

| **RF protein classifier** | **Immune cell phenotype** | **Airway expression status** | **Median airway expression (reporter-corrected intensity)** |  | **Phenotypes in RF classification analysis** |
| --- | --- | --- | --- | --- | --- |
| ABHD17B | B. |  |  |  | B. |
| ACO1 | B. |  |  |  | B.memory |
| AKAP12 | B. |  |  |  | B.naive |
| AKAP13 | B. |  |  |  | B.plasma |
| AKAP2 | B. |  |  |  | Basophil |
| ALDH2 | B. | detected in airway proteome | 145708.5 |  | DC |
| ALOX5 | B. | detected in airway proteome | 9073.1 |  | Eosinophil |
| ARGLU1 | B. |  |  |  | mDC |
| ARHGAP1 | B. | detected in airway proteome | 31323.1 |  | MO |
| ARHGAP24 | B. |  |  |  | MO.classical |
| ARL8A | B. |  |  |  | MO.intermediate |
| ATP6V1H | B. | detected in airway proteome | 12911.4 |  | MO.nonclassical |
| BANK1 | B. |  |  |  | mTregs |
| BASP1 | B. | detected in airway proteome | 5452.7 |  | Neutrophil |
| BLK | B. |  |  |  | NK. |
| BLNK | B. |  |  |  | NK.bright |
| BTK | B. |  |  |  | NK.dim |
| CAMK2D | B. | detected in airway proteome | 41672.8 |  | nTregs |
| CAPZA2 | B. | detected in airway proteome | 49645.1 |  | pDC |
| CAT | B. | detected in airway proteome | 110621.3 |  | T4 |
| CD19 | B. |  |  |  | T4.CM |
| CD22 | B. |  |  |  | T4.EM |
| CD37 | B. |  |  |  | T4.EMRA |
| CD72 | B. |  |  |  | T4.naive |
| CD79A | B. |  |  |  | T8 |
| CD79B | B. |  |  |  | T8.CM |
| CD82 | B. |  |  |  | T8.EM |
| CD9 | B. | detected in airway proteome | 19143.4 |  | T8.EMRA |
| CDAN1 | B. |  |  |  | T8.naive |
| CENPC | B. |  |  |  | Th1 |
| CLEC3B | B. |  |  |  | Th17 |
| CLIP2 | B. |  |  |  | Th2 |
| COBLL1 | B. |  |  |  | Treg |
| CYB561A3 | B. |  |  |  |  |
| DAG1 | B. |  |  |  |  |
| DBNL | B. | detected in airway proteome | 75066.3 |  |  |
| EBF1 | B. |  |  |  |  |
| EML4 | B. |  |  |  |  |
| ENPP3 | B. |  |  |  |  |
| FBLN1 | B. |  |  |  |  |
| FBXO33 | B. |  |  |  |  |
| FCER2 | B. |  |  |  |  |
| FCGR2B | B. |  |  |  |  |
| FCRLA | B. |  |  |  |  |
| FRY | B. |  |  |  |  |
| GNG7 | B. |  |  |  |  |
| GPX4 | B. |  |  |  |  |
| HBA1 | B. | detected in airway proteome | 120595.2 |  |  |
| HIP1R | B. |  |  |  |  |
| HMGN5 | B. |  |  |  |  |
| HPGDS | B. |  |  |  |  |
| HSD17B4 | B. | detected in airway proteome | 204591.3 |  |  |
| HSH2D | B. |  |  |  |  |
| IGFBP2 | B. |  |  |  |  |
| IGHE | B. |  |  |  |  |
| IGHM | B. | detected in airway proteome | 145829.0 |  |  |
| IKZF3 | B. |  |  |  |  |
| KDM4B | B. |  |  |  |  |
| KMO | B. |  |  |  |  |
| LANCL1 | B. |  |  |  |  |
| LILRB2 | B. |  |  |  |  |
| LIMD2 | B. |  |  |  |  |
| LMBRD2 | B. |  |  |  |  |
| LNPEP | B. |  |  |  |  |
| LRSAM1 | B. |  |  |  |  |
| LTF | B. | detected in airway proteome | 2673729.5 |  |  |
| MAPK1 | B. | detected in airway proteome | 23068.8 |  |  |
| MATN1 | B. |  |  |  |  |
| MIA | B. |  |  |  |  |
| MICAL1 | B. |  |  |  |  |
| MICAL3 | B. |  |  |  |  |
| MIOS | B. |  |  |  |  |
| MRPL30 | B. |  |  |  |  |
| MS4A1 | B. |  |  |  |  |
| MYO1D | B. | detected in airway proteome | 7623.0 |  |  |
| MYO9B | B. |  |  |  |  |
| NANS | B. | detected in airway proteome | 31653.0 |  |  |
| NCF1 | B. | detected in airway proteome | 30086.9 |  |  |
| NIPSNAP3B | B. |  |  |  |  |
| OSBPL10 | B. |  |  |  |  |
| OSBPL1A | B. |  |  |  |  |
| OTULIN | B. |  |  |  |  |
| PAFAH1B2 | B. | detected in airway proteome | 13201.7 |  |  |
| PAX5 | B. |  |  |  |  |
| PDCD5 | B. |  |  |  |  |
| PDLIM1 | B. | detected in airway proteome | 118059.9 |  |  |
| PDLIM7 | B. |  |  |  |  |
| PLD3 | B. |  |  |  |  |
| PLEKHA2 | B. |  |  |  |  |
| PLEKHF2 | B. |  |  |  |  |
| PLEKHG1 | B. |  |  |  |  |
| POSTN | B. |  |  |  |  |
| POU2AF1 | B. |  |  |  |  |
| RAB30 | B. |  |  |  |  |
| RAB4A | B. |  |  |  |  |
| RALB | B. | detected in airway proteome | 24583.8 |  |  |
| RALGPS2 | B. |  |  |  |  |
| RBBP9 | B. |  |  |  |  |
| RGN | B. |  |  |  |  |
| RGPD1 | B. |  |  |  |  |
| SEMA7A | B. |  |  |  |  |
| SERPINB1 | B. | detected in airway proteome | 602949.7 |  |  |
| SERPINF1 | B. |  |  |  |  |
| SERPINF2 | B. | detected in airway proteome | 17552.2 |  |  |
| SGPP1 | B. |  |  |  |  |
| SNX18 | B. |  |  |  |  |
| SNX29P2 | B. |  |  |  |  |
| SP100 | B. | detected in airway proteome | 16310.3 |  |  |
| SP140 | B. |  |  |  |  |
| ST6GAL1 | B. | detected in airway proteome | 38047.6 |  |  |
| STAP1 | B. |  |  |  |  |
| STRBP | B. |  |  |  |  |
| SWAP70 | B. |  |  |  |  |
| TANC2 | B. |  |  |  |  |
| TCF3 | B. |  |  |  |  |
| TCL1A | B. |  |  |  |  |
| TDRD7 | B. |  |  |  |  |
| TECPR2 | B. |  |  |  |  |
| TGFBI | B. |  |  |  |  |
| THBS4 | B. |  |  |  |  |
| TLE4 | B. |  |  |  |  |
| TMA7 | B. |  |  |  |  |
| TMPIT | B. |  |  |  |  |
| TPD52 | B. | detected in airway proteome | 24701.4 |  |  |
| TSPAN13 | B. |  |  |  |  |
| TSPAN31 | B. |  |  |  |  |
| TTYH3 | B. |  |  |  |  |
| TXNDC5 | B. | detected in airway proteome | 103227.6 |  |  |
| UBA7 | B. |  |  |  |  |
| VAV3 | B. |  |  |  |  |
| VPREB3 | B. |  |  |  |  |
| VWA5A | B. |  |  |  |  |
| ZC2HC1A | B. |  |  |  |  |
| ZEB1 | B. |  |  |  |  |
| ZHX2 | B. |  |  |  |  |
| ARGLU1 | B.memory |  |  |  |  |
| ARHGAP24 | B.memory |  |  |  |  |
| BANK1 | B.memory |  |  |  |  |
| BLK | B.memory |  |  |  |  |
| C8orf37 | B.memory |  |  |  |  |
| CCDC88C | B.memory |  |  |  |  |
| CD19 | B.memory |  |  |  |  |
| CD1C | B.memory |  |  |  |  |
| CD27 | B.memory |  |  |  |  |
| CD37 | B.memory |  |  |  |  |
| CD79A | B.memory |  |  |  |  |
| CRYZ | B.memory | detected in airway proteome | 34896.4 |  |  |
| DBNL | B.memory | detected in airway proteome | 75066.3 |  |  |
| EBF1 | B.memory |  |  |  |  |
| FCRLA | B.memory |  |  |  |  |
| GPM6A | B.memory |  |  |  |  |
| HIP1R | B.memory |  |  |  |  |
| HMGN3 | B.memory |  |  |  |  |
| IGHM | B.memory | detected in airway proteome | 145829.0 |  |  |
| MAP4K1 | B.memory |  |  |  |  |
| MS4A1 | B.memory |  |  |  |  |
| MYSM1 | B.memory |  |  |  |  |
| PAX5 | B.memory |  |  |  |  |
| PLEKHF2 | B.memory |  |  |  |  |
| PPP1R14A | B.memory |  |  |  |  |
| RAB30 | B.memory |  |  |  |  |
| RALGPS2 | B.memory |  |  |  |  |
| RGPD1 | B.memory |  |  |  |  |
| SP140 | B.memory |  |  |  |  |
| ADK | B.naive | detected in airway proteome | 48022.4 |  |  |
| APOBEC3C | B.naive |  |  |  |  |
| BACH2 | B.naive |  |  |  |  |
| BTLA | B.naive |  |  |  |  |
| CAMK2D | B.naive | detected in airway proteome | 41672.8 |  |  |
| CCL20 | B.naive |  |  |  |  |
| CD19 | B.naive |  |  |  |  |
| CD22 | B.naive |  |  |  |  |
| CD37 | B.naive |  |  |  |  |
| CD72 | B.naive |  |  |  |  |
| CD79A | B.naive |  |  |  |  |
| DENND4A | B.naive |  |  |  |  |
| FCER2 | B.naive |  |  |  |  |
| GLRX3 | B.naive | detected in airway proteome | 33357.1 |  |  |
| GNL1 | B.naive |  |  |  |  |
| GPHN | B.naive |  |  |  |  |
| GTF3C2 | B.naive |  |  |  |  |
| HDAC3 | B.naive |  |  |  |  |
| JAK3 | B.naive |  |  |  |  |
| KAT7 | B.naive |  |  |  |  |
| KMO | B.naive |  |  |  |  |
| MICAL3 | B.naive |  |  |  |  |
| MYCBP2 | B.naive |  |  |  |  |
| MYO1C | B.naive | detected in airway proteome | 23123.4 |  |  |
| NUDCD2 | B.naive |  |  |  |  |
| PATL1 | B.naive |  |  |  |  |
| PAX5 | B.naive |  |  |  |  |
| PIKFYVE | B.naive |  |  |  |  |
| PLEKHA2 | B.naive |  |  |  |  |
| POLDIP3 | B.naive |  |  |  |  |
| QARS | B.naive | detected in airway proteome | 100003.2 |  |  |
| RAB30 | B.naive |  |  |  |  |
| RP9 | B.naive |  |  |  |  |
| SIPA1L3 | B.naive |  |  |  |  |
| SNX29 | B.naive |  |  |  |  |
| SNX29P2 | B.naive |  |  |  |  |
| SNX8 | B.naive |  |  |  |  |
| TCL1A | B.naive |  |  |  |  |
| TNFRSF13B | B.naive |  |  |  |  |
| TNFRSF13C | B.naive |  |  |  |  |
| WDR73 | B.naive |  |  |  |  |
| ZNF652 | B.naive |  |  |  |  |
| ACTN2 | B.plasma |  |  |  |  |
| ALAS1 | B.plasma |  |  |  |  |
| ALDOB | B.plasma |  |  |  |  |
| ANGPTL3 | B.plasma |  |  |  |  |
| ARHGAP11A | B.plasma |  |  |  |  |
| ARID3A | B.plasma |  |  |  |  |
| ASPN | B.plasma |  |  |  |  |
| ASS1 | B.plasma | detected in airway proteome | 34748.2 |  |  |
| B4GALT3 | B.plasma |  |  |  |  |
| C12orf56 | B.plasma |  |  |  |  |
| C1QTNF3 | B.plasma |  |  |  |  |
| CDAN1 | B.plasma |  |  |  |  |
| CLEC11A | B.plasma |  |  |  |  |
| CLEC3B | B.plasma |  |  |  |  |
| COL1A1 | B.plasma | detected in airway proteome | 149886.8 |  |  |
| COL6A1 | B.plasma |  |  |  |  |
| COMP | B.plasma |  |  |  |  |
| CSRP3 | B.plasma |  |  |  |  |
| CWC25 | B.plasma |  |  |  |  |
| DAG1 | B.plasma |  |  |  |  |
| DCD | B.plasma | detected in airway proteome | 17008.6 |  |  |
| EFEMP1 | B.plasma |  |  |  |  |
| FAM122A | B.plasma |  |  |  |  |
| FBLN1 | B.plasma |  |  |  |  |
| FBXO33 | B.plasma |  |  |  |  |
| FMOD | B.plasma |  |  |  |  |
| GIT2 | B.plasma |  |  |  |  |
| GOLM1 | B.plasma | detected in airway proteome | 29708.0 |  |  |
| HBA1 | B.plasma | detected in airway proteome | 120595.2 |  |  |
| HGFAC | B.plasma |  |  |  |  |
| HTRA1 | B.plasma | detected in airway proteome | 13370.8 |  |  |
| IGF2 | B.plasma |  |  |  |  |
| IGFBP2 | B.plasma |  |  |  |  |
| IGFBP3 | B.plasma |  |  |  |  |
| IGFBP4 | B.plasma |  |  |  |  |
| IGFBP5 | B.plasma |  |  |  |  |
| IGHA2 | B.plasma | detected in airway proteome | 18263.5 |  |  |
| IGJ | B.plasma |  |  |  |  |
| KAT6A | B.plasma |  |  |  |  |
| LECT1 | B.plasma |  |  |  |  |
| LMBRD2 | B.plasma |  |  |  |  |
| MAN1A1 | B.plasma |  |  |  |  |
| MAP2 | B.plasma |  |  |  |  |
| MATN1 | B.plasma |  |  |  |  |
| MATN4 | B.plasma |  |  |  |  |
| MFAP5 | B.plasma |  |  |  |  |
| MIA | B.plasma |  |  |  |  |
| MMP15 | B.plasma |  |  |  |  |
| OAF | B.plasma |  |  |  |  |
| PCSK1N | B.plasma |  |  |  |  |
| PLA2G2D | B.plasma |  |  |  |  |
| PLGRKT | B.plasma | detected in airway proteome | 25091.5 |  |  |
| POSTN | B.plasma |  |  |  |  |
| PRSS23 | B.plasma |  |  |  |  |
| PRSS3 | B.plasma |  |  |  |  |
| PRSS35 | B.plasma |  |  |  |  |
| RCN3 | B.plasma |  |  |  |  |
| REST | B.plasma |  |  |  |  |
| RGN | B.plasma |  |  |  |  |
| RORC | B.plasma |  |  |  |  |
| S100A1 | B.plasma |  |  |  |  |
| SERPINF1 | B.plasma |  |  |  |  |
| SERPINF2 | B.plasma | detected in airway proteome | 17552.2 |  |  |
| SMPX | B.plasma |  |  |  |  |
| SPARC | B.plasma |  |  |  |  |
| TAF1C | B.plasma |  |  |  |  |
| TAGLN | B.plasma |  |  |  |  |
| TDRD7 | B.plasma |  |  |  |  |
| TECPR2 | B.plasma |  |  |  |  |
| TGFBI | B.plasma |  |  |  |  |
| TGM3 | B.plasma | detected in airway proteome | 812740.6 |  |  |
| THBS4 | B.plasma |  |  |  |  |
| TRAPPC12 | B.plasma |  |  |  |  |
| TXNDC5 | B.plasma | detected in airway proteome | 103227.6 |  |  |
| ZEB1 | B.plasma |  |  |  |  |
| ABCC1 | Basophil |  |  |  |  |
| ABHD17B | Basophil |  |  |  |  |
| ABHD5 | Basophil | detected in airway proteome | 27880.1 |  |  |
| ACO1 | Basophil |  |  |  |  |
| AKAP12 | Basophil |  |  |  |  |
| ALOX5 | Basophil | detected in airway proteome | 9073.1 |  |  |
| ALS2 | Basophil |  |  |  |  |
| ARSA | Basophil |  |  |  |  |
| ATP7A | Basophil |  |  |  |  |
| BTK | Basophil |  |  |  |  |
| C1orf186 | Basophil |  |  |  |  |
| CA8 | Basophil |  |  |  |  |
| CD151 | Basophil |  |  |  |  |
| CD248 | Basophil |  |  |  |  |
| CD82 | Basophil |  |  |  |  |
| CD9 | Basophil | detected in airway proteome | 19143.4 |  |  |
| CEBPA | Basophil |  |  |  |  |
| CHN2 | Basophil |  |  |  |  |
| CNRIP1 | Basophil |  |  |  |  |
| CPA3 | Basophil |  |  |  |  |
| CRIM1 | Basophil |  |  |  |  |
| CSF2RB | Basophil |  |  |  |  |
| DHRS11 | Basophil |  |  |  |  |
| ENPP3 | Basophil |  |  |  |  |
| FAM101B | Basophil |  |  |  |  |
| FAM110A | Basophil |  |  |  |  |
| FCER1A | Basophil |  |  |  |  |
| FRY | Basophil |  |  |  |  |
| GATA2 | Basophil |  |  |  |  |
| GCSAML | Basophil |  |  |  |  |
| H1F0 | Basophil | detected in airway proteome | 91982.0 |  |  |
| HIST1H1B | Basophil | detected in airway proteome | 149060.6 |  |  |
| HMGN5 | Basophil |  |  |  |  |
| HPGD | Basophil |  |  |  |  |
| HPGDS | Basophil |  |  |  |  |
| HSD17B4 | Basophil | detected in airway proteome | 204591.3 |  |  |
| HYAL3 | Basophil |  |  |  |  |
| IGHE | Basophil |  |  |  |  |
| IMPACT | Basophil |  |  |  |  |
| JAK2 | Basophil |  |  |  |  |
| KLF5 | Basophil |  |  |  |  |
| MFSD6 | Basophil |  |  |  |  |
| MS4A2 | Basophil |  |  |  |  |
| MS4A3 | Basophil |  |  |  |  |
| MYH11 | Basophil |  |  |  |  |
| NFE2 | Basophil |  |  |  |  |
| NHSL2 | Basophil |  |  |  |  |
| NIPSNAP3B | Basophil |  |  |  |  |
| OSBPL1A | Basophil |  |  |  |  |
| PDLIM7 | Basophil |  |  |  |  |
| PLD3 | Basophil |  |  |  |  |
| PPM1H | Basophil |  |  |  |  |
| PRNP | Basophil |  |  |  |  |
| PTGDR2 | Basophil |  |  |  |  |
| RAB37 | Basophil |  |  |  |  |
| RAB4A | Basophil |  |  |  |  |
| RBBP9 | Basophil |  |  |  |  |
| RGS2 | Basophil |  |  |  |  |
| S1PR2 | Basophil |  |  |  |  |
| SGPP1 | Basophil |  |  |  |  |
| SLC18A2 | Basophil |  |  |  |  |
| SLC45A3 | Basophil |  |  |  |  |
| SMPDL3A | Basophil |  |  |  |  |
| SRGAP3 | Basophil |  |  |  |  |
| SSH3 | Basophil |  |  |  |  |
| SYTL3 | Basophil |  |  |  |  |
| TLE4 | Basophil |  |  |  |  |
| TMEM164 | Basophil |  |  |  |  |
| TPSAB1 | Basophil |  |  |  |  |
| TSPAN32 | Basophil |  |  |  |  |
| TTC7B | Basophil |  |  |  |  |
| TTYH3 | Basophil |  |  |  |  |
| UNC13D | Basophil | detected in airway proteome | 9990.5 |  |  |
| VAV3 | Basophil |  |  |  |  |
| VPS13A | Basophil |  |  |  |  |
| VWA5A | Basophil |  |  |  |  |
| YOD1 | Basophil | detected in airway proteome | 26072.6 |  |  |
| ZC2HC1A | Basophil |  |  |  |  |
| ACY3 | DC |  |  |  |  |
| ADA | DC |  |  |  |  |
| AKR1D1 | DC |  |  |  |  |
| ALCAM | DC | detected in airway proteome | 22302.4 |  |  |
| ALDH7A1 | DC | detected in airway proteome | 21503.7 |  |  |
| ALG2 | DC |  |  |  |  |
| ARAP1 | DC |  |  |  |  |
| ATP1B1 | DC | detected in airway proteome | 23211.5 |  |  |
| C12orf45 | DC |  |  |  |  |
| CALCRL | DC |  |  |  |  |
| CCDC50 | DC |  |  |  |  |
| CCDC6 | DC |  |  |  |  |
| CCDC88A | DC |  |  |  |  |
| CD1C | DC |  |  |  |  |
| CD2AP | DC |  |  |  |  |
| CDKN1A | DC |  |  |  |  |
| CKB | DC | detected in airway proteome | 15529.7 |  |  |
| CLASP2 | DC |  |  |  |  |
| CLIC2 | DC |  |  |  |  |
| CMPK2 | DC |  |  |  |  |
| COMT | DC | detected in airway proteome | 43943.8 |  |  |
| CORO1B | DC | detected in airway proteome | 32129.0 |  |  |
| CSRP2 | DC |  |  |  |  |
| CTNNA1 | DC | detected in airway proteome | 24999.0 |  |  |
| CTNND1 | DC | detected in airway proteome | 58039.3 |  |  |
| CXorf21 | DC |  |  |  |  |
| CYP2S1 | DC | detected in airway proteome | 11020.1 |  |  |
| DAB2 | DC |  |  |  |  |
| DCK | DC |  |  |  |  |
| DDAH2 | DC |  |  |  |  |
| DHTKD1 | DC |  |  |  |  |
| DHX58 | DC |  |  |  |  |
| DNAJB4 | DC |  |  |  |  |
| DOCK1 | DC |  |  |  |  |
| DPYSL2 | DC | detected in airway proteome | 10311.7 |  |  |
| DUSP3 | DC |  |  |  |  |
| DUSP5 | DC |  |  |  |  |
| ECE1 | DC |  |  |  |  |
| EIF2AK4 | DC |  |  |  |  |
| EPHA2 | DC |  |  |  |  |
| EPSTI1 | DC |  |  |  |  |
| ETV6 | DC |  |  |  |  |
| FAM213A | DC | detected in airway proteome | 10756.5 |  |  |
| FAM49A | DC |  |  |  |  |
| FCHSD2 | DC |  |  |  |  |
| FEZ2 | DC |  |  |  |  |
| GABARAPL2 | DC |  |  |  |  |
| GCLC | DC |  |  |  |  |
| GPR183 | DC |  |  |  |  |
| GSTM1 | DC | detected in airway proteome | 19223.0 |  |  |
| GSTM2 | DC |  |  |  |  |
| H2AFY2 | DC | detected in airway proteome | 11721.8 |  |  |
| HERC5 | DC |  |  |  |  |
| HIP1 | DC |  |  |  |  |
| HYOU1 | DC | detected in airway proteome | 160839.1 |  |  |
| IDH3A | DC | detected in airway proteome | 63826.7 |  |  |
| IFIH1 | DC |  |  |  |  |
| IFIT1 | DC | detected in airway proteome | 51392.4 |  |  |
| IFIT2 | DC | detected in airway proteome | 13796.5 |  |  |
| IFIT3 | DC |  |  |  |  |
| IRF4 | DC |  |  |  |  |
| IRF7 | DC |  |  |  |  |
| IRF8 | DC |  |  |  |  |
| ISG15 | DC | detected in airway proteome | 37384.4 |  |  |
| ISG20 | DC |  |  |  |  |
| ITGB8 | DC |  |  |  |  |
| KIAA1598 | DC | detected in airway proteome | 13534.8 |  |  |
| KRTCAP2 | DC |  |  |  |  |
| LAP3 | DC | detected in airway proteome | 174126.2 |  |  |
| LGMN | DC |  |  |  |  |
| LILRB2 | DC |  |  |  |  |
| LILRB4 | DC |  |  |  |  |
| LY75 | DC |  |  |  |  |
| MAP1A | DC |  |  |  |  |
| MAP2K6 | DC |  |  |  |  |
| MRC1 | DC |  |  |  |  |
| MTMR12 | DC |  |  |  |  |
| MX2 | DC | detected in airway proteome | 44111.8 |  |  |
| MYO1E | DC | detected in airway proteome | 25874.2 |  |  |
| NCF2 | DC | detected in airway proteome | 80864.6 |  |  |
| NCF4 | DC | detected in airway proteome | 20482.6 |  |  |
| NDRG2 | DC | detected in airway proteome | 34601.9 |  |  |
| NFKB1 | DC | detected in airway proteome | 21136.8 |  |  |
| NFKBIA | DC |  |  |  |  |
| NGLY1 | DC |  |  |  |  |
| NR4A3 | DC |  |  |  |  |
| NUB1 | DC |  |  |  |  |
| OTUD6B | DC |  |  |  |  |
| PACSIN1 | DC |  |  |  |  |
| PAK1 | DC |  |  |  |  |
| PALD1 | DC |  |  |  |  |
| PARP12 | DC |  |  |  |  |
| PFKFB2 | DC |  |  |  |  |
| PIK3R5 | DC |  |  |  |  |
| PLCB3 | DC |  |  |  |  |
| PLCG2 | DC | detected in airway proteome | 18765.2 |  |  |
| PLD4 | DC |  |  |  |  |
| PLEK | DC | detected in airway proteome | 6435.7 |  |  |
| PLXNB2 | DC | detected in airway proteome | 9299.5 |  |  |
| PM20D2 | DC |  |  |  |  |
| PNKP | DC |  |  |  |  |
| PON2 | DC | detected in airway proteome | 4735.4 |  |  |
| PPP1R14B | DC |  |  |  |  |
| PRKAR2B | DC |  |  |  |  |
| PTBP3 | DC |  |  |  |  |
| PTPRE | DC |  |  |  |  |
| PYGL | DC | detected in airway proteome | 57123.5 |  |  |
| REPIN1 | DC |  |  |  |  |
| RPL27 | DC | detected in airway proteome | 115404.9 |  |  |
| RPL3 | DC | detected in airway proteome | 132604.6 |  |  |
| RPL31 | DC | detected in airway proteome | 43773.2 |  |  |
| RPL37A | DC | detected in airway proteome | 32911.0 |  |  |
| RPS6KA4 | DC |  |  |  |  |
| RUFY3 | DC |  |  |  |  |
| S100A3 | DC |  |  |  |  |
| SAMHD1 | DC | detected in airway proteome | 55756.8 |  |  |
| SCML2 | DC |  |  |  |  |
| SCRN1 | DC |  |  |  |  |
| SH2B3 | DC |  |  |  |  |
| SKAP2 | DC |  |  |  |  |
| SMIM5 | DC |  |  |  |  |
| SNCA | DC |  |  |  |  |
| SPECC1 | DC |  |  |  |  |
| SPHK1 | DC |  |  |  |  |
| SRGAP2 | DC |  |  |  |  |
| ST14 | DC |  |  |  |  |
| TAGLN2 | DC | detected in airway proteome | 177934.2 |  |  |
| TARBP1 | DC |  |  |  |  |
| TBC1D13 | DC |  |  |  |  |
| TBC1D4 | DC |  |  |  |  |
| TCP11L1 | DC |  |  |  |  |
| THBD | DC |  |  |  |  |
| TIFAB | DC |  |  |  |  |
| TLR7 | DC |  |  |  |  |
| TMEM206 | DC |  |  |  |  |
| TNS3 | DC |  |  |  |  |
| TRAFD1 | DC |  |  |  |  |
| TRIM22 | DC | detected in airway proteome | 1867.4 |  |  |
| TRIP10 | DC | detected in airway proteome | 11319.0 |  |  |
| TUBB2B | DC |  |  |  |  |
| TUBB6 | DC | detected in airway proteome | 41292.1 |  |  |
| UBE2E2 | DC |  |  |  |  |
| UBE2W | DC |  |  |  |  |
| USP18 | DC |  |  |  |  |
| UVRAG | DC |  |  |  |  |
| VEZF1 | DC |  |  |  |  |
| WDFY4 | DC |  |  |  |  |
| WDR47 | DC |  |  |  |  |
| ZDHHC17 | DC |  |  |  |  |
| ZFP36L1 | DC |  |  |  |  |
| ZNF579 | DC |  |  |  |  |
| ZNF618 | DC |  |  |  |  |
| ZNF672 | DC |  |  |  |  |
| ZNFX1 | DC |  |  |  |  |
| ZNRF2 | DC |  |  |  |  |
| AKR1C1 | Eosinophil | detected in airway proteome | 20065.4 |  |  |
| ALDH3B1 | Eosinophil | detected in airway proteome | 44807.3 |  |  |
| ALOX15 | Eosinophil | detected in airway proteome | 12989.8 |  |  |
| ARSB | Eosinophil |  |  |  |  |
| C19orf35 | Eosinophil |  |  |  |  |
| CAMK1D | Eosinophil |  |  |  |  |
| CCR3 | Eosinophil |  |  |  |  |
| CEBPE | Eosinophil |  |  |  |  |
| CPTP | Eosinophil |  |  |  |  |
| CREG1 | Eosinophil |  |  |  |  |
| DAPK2 | Eosinophil |  |  |  |  |
| DEFA4 | Eosinophil |  |  |  |  |
| DHCR7 | Eosinophil |  |  |  |  |
| EMR1 | Eosinophil |  |  |  |  |
| EPN3 | Eosinophil |  |  |  |  |
| EPX | Eosinophil | detected in airway proteome | 17544.8 |  |  |
| FHL3 | Eosinophil |  |  |  |  |
| G6PD | Eosinophil | detected in airway proteome | 89282.6 |  |  |
| GALC | Eosinophil |  |  |  |  |
| GANAB | Eosinophil | detected in airway proteome | 180081.2 |  |  |
| GAPT | Eosinophil |  |  |  |  |
| GATA1 | Eosinophil |  |  |  |  |
| GSR | Eosinophil | detected in airway proteome | 20483.5 |  |  |
| KEL | Eosinophil |  |  |  |  |
| LGALS12 | Eosinophil |  |  |  |  |
| LPCAT2 | Eosinophil |  |  |  |  |
| LTC4S | Eosinophil |  |  |  |  |
| MYL12B | Eosinophil |  |  |  |  |
| NCF1 | Eosinophil | detected in airway proteome | 30086.9 |  |  |
| PADI2 | Eosinophil | detected in airway proteome | 45551.4 |  |  |
| PIK3R6 | Eosinophil |  |  |  |  |
| PPP1R1B | Eosinophil |  |  |  |  |
| PRG2 | Eosinophil | detected in airway proteome | 26501.4 |  |  |
| PRG3 | Eosinophil | detected in airway proteome | 8400.7 |  |  |
| PRSS33 | Eosinophil |  |  |  |  |
| PSTPIP2 | Eosinophil |  |  |  |  |
| RGS18 | Eosinophil |  |  |  |  |
| RNASE2 | Eosinophil | detected in airway proteome | 18499.5 |  |  |
| RNASE3 | Eosinophil | detected in airway proteome | 76108.9 |  |  |
| RPS6KA2 | Eosinophil |  |  |  |  |
| SIGLEC8 | Eosinophil |  |  |  |  |
| SLC25A51 | Eosinophil |  |  |  |  |
| SMPD3 | Eosinophil |  |  |  |  |
| SPNS3 | Eosinophil |  |  |  |  |
| ST3GAL2 | Eosinophil |  |  |  |  |
| ST5 | Eosinophil |  |  |  |  |
| STXBP5 | Eosinophil |  |  |  |  |
| TM7SF3 | Eosinophil |  |  |  |  |
| WIPI1 | Eosinophil |  |  |  |  |
| ABHD15 | mDC |  |  |  |  |
| ACSF2 | mDC | detected in airway proteome | 14823.8 |  |  |
| AGPAT9 | mDC |  |  |  |  |
| AIM1 | mDC | detected in airway proteome | 76153.7 |  |  |
| ALCAM | mDC | detected in airway proteome | 22302.4 |  |  |
| ALDH7A1 | mDC | detected in airway proteome | 21503.7 |  |  |
| ATP1B1 | mDC | detected in airway proteome | 23211.5 |  |  |
| BCL3 | mDC |  |  |  |  |
| CCDC6 | mDC |  |  |  |  |
| CCDC88A | mDC |  |  |  |  |
| CCR7 | mDC |  |  |  |  |
| CD1C | mDC |  |  |  |  |
| CDKN1A | mDC |  |  |  |  |
| CLEC10A | mDC |  |  |  |  |
| CLIC2 | mDC |  |  |  |  |
| CPD | mDC |  |  |  |  |
| CRLF2 | mDC |  |  |  |  |
| CSF2RA | mDC |  |  |  |  |
| CTNNA1 | mDC | detected in airway proteome | 24999.0 |  |  |
| CTNND1 | mDC | detected in airway proteome | 58039.3 |  |  |
| CYP2S1 | mDC | detected in airway proteome | 11020.1 |  |  |
| DEPTOR | mDC |  |  |  |  |
| DTNA | mDC |  |  |  |  |
| ETV3 | mDC |  |  |  |  |
| FAM49A | mDC |  |  |  |  |
| FECH | mDC | detected in airway proteome | 16734.7 |  |  |
| FSCN1 | mDC | detected in airway proteome | 37645.7 |  |  |
| GABARAPL2 | mDC |  |  |  |  |
| GCLC | mDC |  |  |  |  |
| GPR183 | mDC |  |  |  |  |
| H2AFY2 | mDC | detected in airway proteome | 11721.8 |  |  |
| HAAO | mDC |  |  |  |  |
| HMGCS1 | mDC | detected in airway proteome | 27684.4 |  |  |
| HOMER2 | mDC |  |  |  |  |
| IL12B | mDC |  |  |  |  |
| il15ra | mDC |  |  |  |  |
| IL18 | mDC | detected in airway proteome | 24745.9 |  |  |
| IL6 | mDC |  |  |  |  |
| ITGB8 | mDC |  |  |  |  |
| JUN | mDC |  |  |  |  |
| LGALS2 | mDC |  |  |  |  |
| LY75 | mDC |  |  |  |  |
| MAFF | mDC |  |  |  |  |
| MAPKBP1 | mDC |  |  |  |  |
| MKL1 | mDC |  |  |  |  |
| MRC1 | mDC |  |  |  |  |
| NAV1 | mDC |  |  |  |  |
| NDRG2 | mDC | detected in airway proteome | 34601.9 |  |  |
| NFKB1 | mDC | detected in airway proteome | 21136.8 |  |  |
| NFKBIA | mDC |  |  |  |  |
| NR4A3 | mDC |  |  |  |  |
| NRP2 | mDC |  |  |  |  |
| NSMAF | mDC |  |  |  |  |
| NUB1 | mDC |  |  |  |  |
| PAK1 | mDC |  |  |  |  |
| PKIB | mDC |  |  |  |  |
| PLEKHB2 | mDC |  |  |  |  |
| PM20D2 | mDC |  |  |  |  |
| PON2 | mDC | detected in airway proteome | 4735.4 |  |  |
| RASSF4 | mDC |  |  |  |  |
| RCAN1 | mDC |  |  |  |  |
| RCN2 | mDC | detected in airway proteome | 33305.7 |  |  |
| RPS6KA3 | mDC |  |  |  |  |
| RTN1 | mDC |  |  |  |  |
| RUFY3 | mDC |  |  |  |  |
| SPATS2L | mDC | detected in airway proteome |  |  |  |
| SPECC1 | mDC |  |  |  |  |
| SQSTM1 | mDC |  |  |  |  |
| STK38L | mDC |  |  |  |  |
| TET2 | mDC |  |  |  |  |
| TNFAIP2 | mDC |  |  |  |  |
| TRIO | mDC |  |  |  |  |
| TRIP10 | mDC | detected in airway proteome | 11319.0 |  |  |
| TRPS1 | mDC |  |  |  |  |
| ZNF579 | mDC |  |  |  |  |
| ZNF672 | mDC |  |  |  |  |
| ACSL1 | MO | detected in airway proteome | 68941.9 |  |  |
| AGTRAP | MO |  |  |  |  |
| AHNAK | MO | detected in airway proteome | 413184.4 |  |  |
| ALDH1A1 | MO | detected in airway proteome | 68038.3 |  |  |
| ALDH3A2 | MO | detected in airway proteome | 49187.2 |  |  |
| ANXA2 | MO | detected in airway proteome | 1356986.0 |  |  |
| ANXA5 | MO | detected in airway proteome | 323252.6 |  |  |
| APOBEC3A | MO | detected in airway proteome | 21532.9 |  |  |
| ARHGEF10L | MO |  |  |  |  |
| ATP5I | MO | detected in airway proteome | 38562.1 |  |  |
| ATP6AP1 | MO |  |  |  |  |
| ATP6V0D2 | MO |  |  |  |  |
| ATP6V1B2 | MO | detected in airway proteome | 87153.8 |  |  |
| ATP6V1G1 | MO | detected in airway proteome | 27236.9 |  |  |
| BACH1 | MO |  |  |  |  |
| BLVRB | MO | detected in airway proteome | 21369.6 |  |  |
| C6orf120 | MO |  |  |  |  |
| CD14 | MO | detected in airway proteome | 13810.2 |  |  |
| CD300E | MO |  |  |  |  |
| CD36 | MO |  |  |  |  |
| CDC6 | MO |  |  |  |  |
| CES1 | MO | detected in airway proteome | 102350.7 |  |  |
| CKAP4 | MO | detected in airway proteome | 61515.6 |  |  |
| CLPB | MO |  |  |  |  |
| COLGALT1 | MO | detected in airway proteome | 2477.9 |  |  |
| COMMD10 | MO |  |  |  |  |
| CRAT | MO | detected in airway proteome | 39625.1 |  |  |
| CTSL | MO |  |  |  |  |
| CYC1 | MO | detected in airway proteome | 45848.9 |  |  |
| CYP27A1 | MO |  |  |  |  |
| DIAPH2 | MO |  |  |  |  |
| DPYD | MO |  |  |  |  |
| DSCR3 | MO |  |  |  |  |
| F13A1 | MO |  |  |  |  |
| FAM45A | MO |  |  |  |  |
| FCGR1A | MO |  |  |  |  |
| GAA | MO |  |  |  |  |
| GHDC | MO |  |  |  |  |
| HADHA | MO | detected in airway proteome | 335279.5 |  |  |
| hCG_14925 | MO |  |  |  |  |
| HEBP1 | MO |  |  |  |  |
| HMOX1 | MO | detected in airway proteome | 28938.0 |  |  |
| HNMT | MO | detected in airway proteome | 8498.2 |  |  |
| IDH1 | MO | detected in airway proteome | 117112.5 |  |  |
| IFI30 | MO | detected in airway proteome | 9511.2 |  |  |
| IRAK3 | MO |  |  |  |  |
| LAMTOR1 | MO |  |  |  |  |
| LDHD | MO |  |  |  |  |
| LGALS1 | MO | detected in airway proteome | 34563.2 |  |  |
| LRP1 | MO |  |  |  |  |
| LRPAP1 | MO |  |  |  |  |
| LRRC25 | MO |  |  |  |  |
| MFN2 | MO |  |  |  |  |
| MGAT1 | MO |  |  |  |  |
| NAIP | MO |  |  |  |  |
| NAPRT | MO | detected in airway proteome | 28659.2 |  |  |
| NCEH1 | MO |  |  |  |  |
| NDUFB8 | MO | detected in airway proteome | 14007.1 |  |  |
| NPL | MO |  |  |  |  |
| PICALM | MO |  |  |  |  |
| PLEKHA4 | MO |  |  |  |  |
| PLOD1 | MO | detected in airway proteome | 23358.3 |  |  |
| PNKD | MO |  |  |  |  |
| POR | MO | detected in airway proteome | 72721.3 |  |  |
| PYCARD | MO |  |  |  |  |
| RAB5A | MO | detected in airway proteome | 18249.2 |  |  |
| RHOT1 | MO |  |  |  |  |
| RIT1 | MO | detected in airway proteome | 5188.0 |  |  |
| S100A4 | MO | detected in airway proteome | 61614.6 |  |  |
| S100A6 | MO | detected in airway proteome | 70082.5 |  |  |
| SERPINB8 | MO | detected in airway proteome |  |  |  |
| SFXN3 | MO | detected in airway proteome | 8107.9 |  |  |
| SGPL1 | MO |  |  |  |  |
| SLC12A9 | MO |  |  |  |  |
| SLC27A3 | MO |  |  |  |  |
| SLC31A1 | MO |  |  |  |  |
| SPTLC1 | MO |  |  |  |  |
| STAB1 | MO |  |  |  |  |
| STX11 | MO |  |  |  |  |
| SULT1A1 | MO |  |  |  |  |
| SULT1A4 | MO |  |  |  |  |
| SYNE3 | MO |  |  |  |  |
| TBC1D2 | MO |  |  |  |  |
| TLR2 | MO |  |  |  |  |
| TLR8 | MO |  |  |  |  |
| TM9SF4 | MO | detected in airway proteome | 11889.1 |  |  |
| TOLLIP | MO | detected in airway proteome | 35011.8 |  |  |
| TSPO | MO |  |  |  |  |
| TYMP | MO | detected in airway proteome | 27710.1 |  |  |
| UQCRC1 | MO | detected in airway proteome | 176655.1 |  |  |
| UQCRC2 | MO | detected in airway proteome | 174732.5 |  |  |
| UQCRFS1 | MO | detected in airway proteome | 28991.3 |  |  |
| ACTG1 | MO.classical | detected in airway proteome | 9420.3 |  |  |
| AGA | MO.classical |  |  |  |  |
| AGTRAP | MO.classical |  |  |  |  |
| ATG13 | MO.classical |  |  |  |  |
| ATG2A | MO.classical |  |  |  |  |
| CCR1 | MO.classical |  |  |  |  |
| CD14 | MO.classical | detected in airway proteome | 13810.2 |  |  |
| CD163 | MO.classical |  |  |  |  |
| CD36 | MO.classical |  |  |  |  |
| CD37 | MO.classical |  |  |  |  |
| CDCA7L | MO.classical |  |  |  |  |
| CEBPD | MO.classical |  |  |  |  |
| CEP152 | MO.classical |  |  |  |  |
| CES1 | MO.classical | detected in airway proteome | 102350.7 |  |  |
| CKAP4 | MO.classical | detected in airway proteome | 61515.6 |  |  |
| CLEC4E | MO.classical |  |  |  |  |
| CPM | MO.classical |  |  |  |  |
| CXCL5 | MO.classical |  |  |  |  |
| CXCL8 | MO.classical |  |  |  |  |
| CYP27A1 | MO.classical |  |  |  |  |
| DYNLT1 | MO.classical |  |  |  |  |
| ELOVL7 | MO.classical |  |  |  |  |
| FABP4 | MO.classical |  |  |  |  |
| FAM129B | MO.classical | detected in airway proteome | 166631.5 |  |  |
| FAM160A2 | MO.classical |  |  |  |  |
| FCAR | MO.classical | detected in airway proteome | 36796.0 |  |  |
| FCGR1A | MO.classical |  |  |  |  |
| FPR1 | MO.classical |  |  |  |  |
| FPR2 | MO.classical |  |  |  |  |
| FTH1 | MO.classical | detected in airway proteome | 26204.8 |  |  |
| GABARAP | MO.classical |  |  |  |  |
| GNAQ | MO.classical | detected in airway proteome | 22611.7 |  |  |
| GUSB | MO.classical |  |  |  |  |
| HLTF | MO.classical |  |  |  |  |
| IL1B | MO.classical |  |  |  |  |
| IQGAP3 | MO.classical |  |  |  |  |
| ITGA5 | MO.classical |  |  |  |  |
| LACC1 | MO.classical |  |  |  |  |
| LAMTOR4 | MO.classical |  |  |  |  |
| LTB4R | MO.classical |  |  |  |  |
| LYZ | MO.classical | detected in airway proteome | 204290.5 |  |  |
| METTL7B | MO.classical |  |  |  |  |
| MPP1 | MO.classical |  |  |  |  |
| MTFMT | MO.classical |  |  |  |  |
| NCSTN | MO.classical | detected in airway proteome | 22189.9 |  |  |
| NINJ1 | MO.classical |  |  |  |  |
| OBFC1 | MO.classical |  |  |  |  |
| PAM | MO.classical |  |  |  |  |
| PLAC8 | MO.classical |  |  |  |  |
| RAD9A | MO.classical |  |  |  |  |
| RALGAPA1 | MO.classical |  |  |  |  |
| RBM47 | MO.classical | detected in airway proteome | 31763.5 |  |  |
| RIT1 | MO.classical | detected in airway proteome | 5188.0 |  |  |
| RPL6 | MO.classical | detected in airway proteome | 114204.5 |  |  |
| SERPINB1 | MO.classical | detected in airway proteome | 602949.7 |  |  |
| SERPINB13 | MO.classical | detected in airway proteome | 40249.1 |  |  |
| SERPINB2 | MO.classical | detected in airway proteome | 37709.6 |  |  |
| SLC44A1 | MO.classical |  |  |  |  |
| SLC7A6 | MO.classical |  |  |  |  |
| STEAP3 | MO.classical |  |  |  |  |
| SUN1 | MO.classical |  |  |  |  |
| TMEM50A | MO.classical |  |  |  |  |
| VCAN | MO.classical |  |  |  |  |
| ZNF185 | MO.classical | detected in airway proteome | 152509.2 |  |  |
| ACTR1A | MO.intermediate | detected in airway proteome | 90654.0 |  |  |
| ADAP1 | MO.intermediate |  |  |  |  |
| AKAP13 | MO.intermediate |  |  |  |  |
| ALDH1A1 | MO.intermediate | detected in airway proteome | 68038.3 |  |  |
| AMY1A | MO.intermediate | detected in airway proteome | 730462.6 |  |  |
| AP3B1 | MO.intermediate |  |  |  |  |
| APOBEC3A | MO.intermediate | detected in airway proteome | 21532.9 |  |  |
| ASNA1 | MO.intermediate | detected in airway proteome | 10387.2 |  |  |
| ATP6V1B2 | MO.intermediate | detected in airway proteome | 87153.8 |  |  |
| ATP6V1H | MO.intermediate | detected in airway proteome | 12911.4 |  |  |
| BLVRB | MO.intermediate | detected in airway proteome | 21369.6 |  |  |
| BRD3 | MO.intermediate |  |  |  |  |
| CAPN2 | MO.intermediate | detected in airway proteome | 23004.9 |  |  |
| CASP1 | MO.intermediate |  |  |  |  |
| CCND2 | MO.intermediate |  |  |  |  |
| CHCHD5 | MO.intermediate |  |  |  |  |
| CLINT1 | MO.intermediate |  |  |  |  |
| CLTC | MO.intermediate | detected in airway proteome | 279320.4 |  |  |
| COMMD2 | MO.intermediate |  |  |  |  |
| CYFIP1 | MO.intermediate |  |  |  |  |
| DERA | MO.intermediate | detected in airway proteome | 25705.0 |  |  |
| DNAJB12 | MO.intermediate |  |  |  |  |
| DNM1L | MO.intermediate |  |  |  |  |
| DYNC1LI2 | MO.intermediate | detected in airway proteome | 14676.1 |  |  |
| F13A1 | MO.intermediate |  |  |  |  |
| FKBP5 | MO.intermediate |  |  |  |  |
| FLNA | MO.intermediate | detected in airway proteome | 345214.6 |  |  |
| GALK1 | MO.intermediate |  |  |  |  |
| GIT1 | MO.intermediate |  |  |  |  |
| GPANK1 | MO.intermediate |  |  |  |  |
| GPX1 | MO.intermediate | detected in airway proteome | 48961.2 |  |  |
| HECTD1 | MO.intermediate |  |  |  |  |
| HIST2H3PS2 | MO.intermediate | detected in airway proteome | 41689.3 |  |  |
| IDH1 | MO.intermediate | detected in airway proteome | 117112.5 |  |  |
| IER2 | MO.intermediate |  |  |  |  |
| ILK | MO.intermediate | detected in airway proteome | 8889.2 |  |  |
| INPPL1 | MO.intermediate |  |  |  |  |
| IRAK4 | MO.intermediate |  |  |  |  |
| KIF13B | MO.intermediate |  |  |  |  |
| MAP3K11 | MO.intermediate |  |  |  |  |
| MEFV | MO.intermediate |  |  |  |  |
| MSN | MO.intermediate | detected in airway proteome | 133174.0 |  |  |
| MTOR | MO.intermediate |  |  |  |  |
| NAGK | MO.intermediate | detected in airway proteome | 54775.8 |  |  |
| NDST2 | MO.intermediate |  |  |  |  |
| NEDD4L | MO.intermediate |  |  |  |  |
| NR3C1 | MO.intermediate |  |  |  |  |
| NUBP1 | MO.intermediate |  |  |  |  |
| NXT2 | MO.intermediate |  |  |  |  |
| OSBPL11 | MO.intermediate |  |  |  |  |
| OSBPL9 | MO.intermediate |  |  |  |  |
| PDDC1 | MO.intermediate |  |  |  |  |
| PFKL | MO.intermediate | detected in airway proteome | 38940.8 |  |  |
| PGLS | MO.intermediate | detected in airway proteome | 32820.8 |  |  |
| PICALM | MO.intermediate |  |  |  |  |
| PID1 | MO.intermediate |  |  |  |  |
| PLIN3 | MO.intermediate | detected in airway proteome | 52927.9 |  |  |
| PREP | MO.intermediate |  |  |  |  |
| PRKCB | MO.intermediate | detected in airway proteome | 31610.2 |  |  |
| RABGAP1 | MO.intermediate |  |  |  |  |
| RIPK3 | MO.intermediate |  |  |  |  |
| RNH1 | MO.intermediate | detected in airway proteome | 90905.9 |  |  |
| RPL18A | MO.intermediate | detected in airway proteome | 91049.0 |  |  |
| S100A6 | MO.intermediate | detected in airway proteome | 70082.5 |  |  |
| SCYL2 | MO.intermediate |  |  |  |  |
| SH3BGRL | MO.intermediate | detected in airway proteome | 22100.2 |  |  |
| SIRPB2 | MO.intermediate |  |  |  |  |
| SMARCA4 | MO.intermediate |  |  |  |  |
| SPR | MO.intermediate | detected in airway proteome | 10103.4 |  |  |
| STAG2 | MO.intermediate |  |  |  |  |
| SUCLA2 | MO.intermediate | detected in airway proteome | 52101.1 |  |  |
| SULT1A4 | MO.intermediate |  |  |  |  |
| TAF6 | MO.intermediate |  |  |  |  |
| TARBP2 | MO.intermediate |  |  |  |  |
| TBC1D2 | MO.intermediate |  |  |  |  |
| TIMM50 | MO.intermediate |  |  |  |  |
| TLN1 | MO.intermediate | detected in airway proteome | 36949.8 |  |  |
| TLR9 | MO.intermediate |  |  |  |  |
| TMEM231 | MO.intermediate |  |  |  |  |
| TMPO | MO.intermediate |  |  |  |  |
| TNFRSF21 | MO.intermediate |  |  |  |  |
| TPMT | MO.intermediate |  |  |  |  |
| TRAPPC5 | MO.intermediate |  |  |  |  |
| TRIM25 | MO.intermediate | detected in airway proteome | 10335.7 |  |  |
| USP8 | MO.intermediate |  |  |  |  |
| VPS18 | MO.intermediate |  |  |  |  |
| VPS26A | MO.intermediate | detected in airway proteome | 41801.7 |  |  |
| VPS53 | MO.intermediate |  |  |  |  |
| ZZEF1 | MO.intermediate |  |  |  |  |
| ABCD1 | MO.nonclassical |  |  |  |  |
| ABCD3 | MO.nonclassical | detected in airway proteome | 25350.8 |  |  |
| ABHD16A | MO.nonclassical |  |  |  |  |
| ADCY7 | MO.nonclassical |  |  |  |  |
| ADRBK2 | MO.nonclassical |  |  |  |  |
| AGPAT3 | MO.nonclassical |  |  |  |  |
| ALDH3A2 | MO.nonclassical | detected in airway proteome | 49187.2 |  |  |
| AREL1 | MO.nonclassical |  |  |  |  |
| ARL6IP5 | MO.nonclassical |  |  |  |  |
| ARRB1 | MO.nonclassical |  |  |  |  |
| ATP2B1 | MO.nonclassical |  |  |  |  |
| ATP5O | MO.nonclassical | detected in airway proteome | 135605.2 |  |  |
| ATP6V0D2 | MO.nonclassical |  |  |  |  |
| BIN3 | MO.nonclassical |  |  |  |  |
| C6orf89 | MO.nonclassical |  |  |  |  |
| C9orf89 | MO.nonclassical |  |  |  |  |
| CEBPB | MO.nonclassical |  |  |  |  |
| CERK | MO.nonclassical |  |  |  |  |
| CHST2 | MO.nonclassical |  |  |  |  |
| CHST7 | MO.nonclassical |  |  |  |  |
| CLN6 | MO.nonclassical |  |  |  |  |
| CNIH4 | MO.nonclassical |  |  |  |  |
| CX3CR1 | MO.nonclassical |  |  |  |  |
| DAPK1 | MO.nonclassical |  |  |  |  |
| EMC10 | MO.nonclassical | detected in airway proteome | 7351.8 |  |  |
| EMC3 | MO.nonclassical |  |  |  |  |
| ERGIC1 | MO.nonclassical |  |  |  |  |
| FAR1 | MO.nonclassical |  |  |  |  |
| GIMAP2 | MO.nonclassical |  |  |  |  |
| GNGT2 | MO.nonclassical |  |  |  |  |
| GOLT1B | MO.nonclassical |  |  |  |  |
| HADHA | MO.nonclassical | detected in airway proteome | 335279.5 |  |  |
| hCG_14925 | MO.nonclassical |  |  |  |  |
| HK1 | MO.nonclassical | detected in airway proteome | 147880.9 |  |  |
| HMOX1 | MO.nonclassical | detected in airway proteome | 28938.0 |  |  |
| ITGAX | MO.nonclassical | detected in airway proteome | 10958.5 |  |  |
| ITGB1 | MO.nonclassical |  |  |  |  |
| KDELR1 | MO.nonclassical |  |  |  |  |
| LACTB | MO.nonclassical |  |  |  |  |
| LILRB2 | MO.nonclassical |  |  |  |  |
| LIPA | MO.nonclassical |  |  |  |  |
| LPCAT3 | MO.nonclassical |  |  |  |  |
| LRRC25 | MO.nonclassical |  |  |  |  |
| LY6E | MO.nonclassical |  |  |  |  |
| LYN | MO.nonclassical | detected in airway proteome | 10071.8 |  |  |
| LYPD2 | MO.nonclassical | detected in airway proteome | 6115.5 |  |  |
| MCOLN1 | MO.nonclassical |  |  |  |  |
| METTL7A | MO.nonclassical |  |  |  |  |
| MFSD10 | MO.nonclassical |  |  |  |  |
| MFSD12 | MO.nonclassical |  |  |  |  |
| MPDU1 | MO.nonclassical |  |  |  |  |
| MTCH1 | MO.nonclassical |  |  |  |  |
| MTSS1 | MO.nonclassical |  |  |  |  |
| NDUFA13 | MO.nonclassical | detected in airway proteome | 15809.1 |  |  |
| NDUFB1 | MO.nonclassical | detected in airway proteome | 9039.5 |  |  |
| PAPSS2 | MO.nonclassical | detected in airway proteome |  |  |  |
| PLOD3 | MO.nonclassical |  |  |  |  |
| PNPLA6 | MO.nonclassical |  |  |  |  |
| POFUT1 | MO.nonclassical | detected in airway proteome | 20581.9 |  |  |
| PTGS1 | MO.nonclassical |  |  |  |  |
| PTPLAD2 | MO.nonclassical |  |  |  |  |
| RAB39A | MO.nonclassical | detected in airway proteome | 19074.5 |  |  |
| RAB42 | MO.nonclassical |  |  |  |  |
| RAB5A | MO.nonclassical | detected in airway proteome | 18249.2 |  |  |
| RAPH1 | MO.nonclassical |  |  |  |  |
| RHOB | MO.nonclassical |  |  |  |  |
| RHOC | MO.nonclassical |  |  |  |  |
| RIN1 | MO.nonclassical |  |  |  |  |
| RRAS | MO.nonclassical |  |  |  |  |
| RXRA | MO.nonclassical |  |  |  |  |
| SAMM50 | MO.nonclassical | detected in airway proteome |  |  |  |
| SCCPDH | MO.nonclassical | detected in airway proteome | 18099.7 |  |  |
| SGPL1 | MO.nonclassical |  |  |  |  |
| SH2B2 | MO.nonclassical |  |  |  |  |
| SIGLEC10 | MO.nonclassical |  |  |  |  |
| SLC25A13 | MO.nonclassical | detected in airway proteome | 13559.7 |  |  |
| SLC25A35 | MO.nonclassical |  |  |  |  |
| SLC25A40 | MO.nonclassical |  |  |  |  |
| SLC27A3 | MO.nonclassical |  |  |  |  |
| SLC27A4 | MO.nonclassical |  |  |  |  |
| SLC2A6 | MO.nonclassical |  |  |  |  |
| SLC8B1 | MO.nonclassical |  |  |  |  |
| SPI1 | MO.nonclassical |  |  |  |  |
| SRC | MO.nonclassical |  |  |  |  |
| STRIP1 | MO.nonclassical |  |  |  |  |
| TBC1D8 | MO.nonclassical |  |  |  |  |
| TLR1 | MO.nonclassical |  |  |  |  |
| TMEM259 | MO.nonclassical |  |  |  |  |
| TOR4A | MO.nonclassical |  |  |  |  |
| TPPP3 | MO.nonclassical | detected in airway proteome | 97036.0 |  |  |
| TRIM14 | MO.nonclassical |  |  |  |  |
| UBASH3B | MO.nonclassical |  |  |  |  |
| UNC119 | MO.nonclassical |  |  |  |  |
| VDAC2 | MO.nonclassical | detected in airway proteome | 333048.0 |  |  |
| WDR11 | MO.nonclassical |  |  |  |  |
| ZBTB7A | MO.nonclassical |  |  |  |  |
| ABCC1 | mTregs |  |  |  |  |
| ATRX | mTregs |  |  |  |  |
| CCR10 | mTregs |  |  |  |  |
| CLPTM1L | mTregs |  |  |  |  |
| EPB41 | mTregs |  |  |  |  |
| ERBB2IP | mTregs |  |  |  |  |
| FAS | mTregs |  |  |  |  |
| FOXP3 | mTregs |  |  |  |  |
| GLCE | mTregs |  |  |  |  |
| INTS10 | mTregs |  |  |  |  |
| LAX1 | mTregs |  |  |  |  |
| LPAR2 | mTregs |  |  |  |  |
| MCRS1 | mTregs |  |  |  |  |
| MEOX1 | mTregs |  |  |  |  |
| NIPBL | mTregs | detected in airway proteome | 37022.4 |  |  |
| NTHL1 | mTregs |  |  |  |  |
| OCIAD2 | mTregs |  |  |  |  |
| PGM2L1 | mTregs |  |  |  |  |
| PI16 | mTregs |  |  |  |  |
| POU2F1 | mTregs |  |  |  |  |
| RBBP6 | mTregs |  |  |  |  |
| SLC35D1 | mTregs |  |  |  |  |
| TAF4 | mTregs |  |  |  |  |
| TERF2 | mTregs |  |  |  |  |
| TRIM16 | mTregs | detected in airway proteome | 43295.9 |  |  |
| ZC2HC1A | mTregs |  |  |  |  |
| ACOX1 | Neutrophil | detected in airway proteome | 46948.2 |  |  |
| ADAM10 | Neutrophil |  |  |  |  |
| ADAMTS13 | Neutrophil |  |  |  |  |
| ADD3 | Neutrophil |  |  |  |  |
| AK6 | Neutrophil |  |  |  |  |
| AMFR | Neutrophil |  |  |  |  |
| AMICA1 | Neutrophil |  |  |  |  |
| ANO10 | Neutrophil |  |  |  |  |
| ANO9 | Neutrophil |  |  |  |  |
| ANXA1 | Neutrophil | detected in airway proteome | 1468610.7 |  |  |
| ANXA3 | Neutrophil | detected in airway proteome | 606077.6 |  |  |
| ANXA6 | Neutrophil | detected in airway proteome | 448636.9 |  |  |
| AQP9 | Neutrophil |  |  |  |  |
| ARHGAP30 | Neutrophil |  |  |  |  |
| ARL8A | Neutrophil |  |  |  |  |
| ARMC8 | Neutrophil |  |  |  |  |
| ARRDC1 | Neutrophil |  |  |  |  |
| ASAP1 | Neutrophil |  |  |  |  |
| ATAT1 | Neutrophil |  |  |  |  |
| ATP11B | Neutrophil | detected in airway proteome | 13257.5 |  |  |
| ATP8A1 | Neutrophil |  |  |  |  |
| AZU1 | Neutrophil | detected in airway proteome | 64595.4 |  |  |
| BICD2 | Neutrophil |  |  |  |  |
| BPI | Neutrophil | detected in airway proteome | 104773.6 |  |  |
| CA4 | Neutrophil | detected in airway proteome | 4284.1 |  |  |
| CALM1 | Neutrophil |  |  |  |  |
| CAPZA1 | Neutrophil | detected in airway proteome | 99499.4 |  |  |
| CD46 | Neutrophil |  |  |  |  |
| CD47 | Neutrophil |  |  |  |  |
| CD63 | Neutrophil |  |  |  |  |
| CDA | Neutrophil | detected in airway proteome | 27839.1 |  |  |
| CDC34 | Neutrophil |  |  |  |  |
| CEACAM3 | Neutrophil |  |  |  |  |
| CEACAM8 | Neutrophil | detected in airway proteome | 22764.4 |  |  |
| CENPH | Neutrophil |  |  |  |  |
| CFP | Neutrophil |  |  |  |  |
| CHMP2A | Neutrophil |  |  |  |  |
| CHTF8 | Neutrophil |  |  |  |  |
| CLDND1 | Neutrophil |  |  |  |  |
| CLEC4D | Neutrophil |  |  |  |  |
| COL4A3BP | Neutrophil |  |  |  |  |
| CR1 | Neutrophil | detected in airway proteome | 16970.8 |  |  |
| CRISP3 | Neutrophil | detected in airway proteome | 88620.5 |  |  |
| CSF3R | Neutrophil |  |  |  |  |
| CTSG | Neutrophil | detected in airway proteome | 501844.6 |  |  |
| CXCR1 | Neutrophil |  |  |  |  |
| CYBRD1 | Neutrophil |  |  |  |  |
| CYP4F3 | Neutrophil | detected in airway proteome | 4060.8 |  |  |
| DEF6 | Neutrophil |  |  |  |  |
| DNAJC5 | Neutrophil |  |  |  |  |
| DROSHA | Neutrophil |  |  |  |  |
| ELMO1 | Neutrophil | detected in airway proteome | 24732.5 |  |  |
| ERCC6 | Neutrophil |  |  |  |  |
| ERO1LB | Neutrophil |  |  |  |  |
| EURL | Neutrophil |  |  |  |  |
| EVI2B | Neutrophil |  |  |  |  |
| EXOC6 | Neutrophil |  |  |  |  |
| FGR | Neutrophil | detected in airway proteome | 20204.0 |  |  |
| FLOT1 | Neutrophil | detected in airway proteome | 21307.1 |  |  |
| GBE1 | Neutrophil | detected in airway proteome | 27814.0 |  |  |
| GCA | Neutrophil | detected in airway proteome | 96225.3 |  |  |
| GGCT | Neutrophil | detected in airway proteome | 21012.7 |  |  |
| GLTSCR1L | Neutrophil |  |  |  |  |
| GMIP | Neutrophil |  |  |  |  |
| GNAI3 | Neutrophil | detected in airway proteome | 17512.8 |  |  |
| GNB2 | Neutrophil | detected in airway proteome | 30593.8 |  |  |
| GNB2L1 | Neutrophil | detected in airway proteome | 106695.7 |  |  |
| GNB4 | Neutrophil | detected in airway proteome | 15566.0 |  |  |
| GNG2 | Neutrophil |  |  |  |  |
| GOLGA7 | Neutrophil |  |  |  |  |
| GPN3 | Neutrophil |  |  |  |  |
| GPR15 | Neutrophil |  |  |  |  |
| GPSM3 | Neutrophil |  |  |  |  |
| GRN | Neutrophil | detected in airway proteome | 2717.1 |  |  |
| H6PD | Neutrophil | detected in airway proteome | 9865.9 |  |  |
| HIVEP2 | Neutrophil |  |  |  |  |
| HLTF | Neutrophil |  |  |  |  |
| HOOK3 | Neutrophil |  |  |  |  |
| HP | Neutrophil | detected in airway proteome | 91441.1 |  |  |
| HSP90AA4P | Neutrophil |  |  |  |  |
| IGF2R | Neutrophil |  |  |  |  |
| IP6K1 | Neutrophil |  |  |  |  |
| ITGAM | Neutrophil | detected in airway proteome | 256276.4 |  |  |
| KIR2DL3 | Neutrophil |  |  |  |  |
| KLC4 | Neutrophil |  |  |  |  |
| KXD1 | Neutrophil |  |  |  |  |
| LCLAT1 | Neutrophil |  |  |  |  |
| LCN2 | Neutrophil | detected in airway proteome | 214619.3 |  |  |
| LILRA3 | Neutrophil |  |  |  |  |
| LILRA6 | Neutrophil |  |  |  |  |
| LMAN2 | Neutrophil | detected in airway proteome | 47802.9 |  |  |
| LMBRD1 | Neutrophil |  |  |  |  |
| LMTK2 | Neutrophil |  |  |  |  |
| LNP | Neutrophil |  |  |  |  |
| LUC7L3 | Neutrophil |  |  |  |  |
| MAP2K2 | Neutrophil |  |  |  |  |
| MAP3K1 | Neutrophil |  |  |  |  |
| MAPK14 | Neutrophil | detected in airway proteome | 6076.1 |  |  |
| MCUR1 | Neutrophil |  |  |  |  |
| MFSD9 | Neutrophil |  |  |  |  |
| MICU2 | Neutrophil | detected in airway proteome | 14585.4 |  |  |
| MME | Neutrophil |  |  |  |  |
| MMP9 | Neutrophil | detected in airway proteome | 67160.6 |  |  |
| MSL1 | Neutrophil |  |  |  |  |
| MST4 | Neutrophil |  |  |  |  |
| MYH9 | Neutrophil | detected in airway proteome | 281279.9 |  |  |
| MYL12A | Neutrophil | detected in airway proteome | 139916.6 |  |  |
| MYL6B | Neutrophil |  |  |  |  |
| MYLK | Neutrophil | detected in airway proteome | 9376.0 |  |  |
| MYO18A | Neutrophil |  |  |  |  |
| MYO1F | Neutrophil | detected in airway proteome | 17723.8 |  |  |
| MYO9B | Neutrophil |  |  |  |  |
| NAGPA | Neutrophil |  |  |  |  |
| NDE1 | Neutrophil |  |  |  |  |
| NEDD8 | Neutrophil |  |  |  |  |
| NKIRAS2 | Neutrophil |  |  |  |  |
| NMRAL1 | Neutrophil |  |  |  |  |
| NQO2 | Neutrophil |  |  |  |  |
| NT5DC3 | Neutrophil |  |  |  |  |
| OLFM4 | Neutrophil | detected in airway proteome | 44952.9 |  |  |
| ORM1 | Neutrophil | detected in airway proteome | 51439.7 |  |  |
| P4HTM | Neutrophil |  |  |  |  |
| PACSIN2 | Neutrophil |  |  |  |  |
| PCBD2 | Neutrophil |  |  |  |  |
| PDE6H | Neutrophil |  |  |  |  |
| PFN1 | Neutrophil | detected in airway proteome | 219106.2 |  |  |
| PGD | Neutrophil | detected in airway proteome | 213588.9 |  |  |
| PGLYRP1 | Neutrophil | detected in airway proteome | 25473.9 |  |  |
| PHF12 | Neutrophil |  |  |  |  |
| PHKG2 | Neutrophil |  |  |  |  |
| PKLR | Neutrophil |  |  |  |  |
| PLAA | Neutrophil |  |  |  |  |
| POLM | Neutrophil |  |  |  |  |
| PPM1B | Neutrophil |  |  |  |  |
| PPP1R12A | Neutrophil |  |  |  |  |
| PPP1R18 | Neutrophil |  |  |  |  |
| PPP2CA | Neutrophil | detected in airway proteome | 20454.9 |  |  |
| PPP3CA | Neutrophil |  |  |  |  |
| PRKAR1A | Neutrophil | detected in airway proteome | 16035.1 |  |  |
| PRKCSH | Neutrophil | detected in airway proteome | 33786.6 |  |  |
| PSMB9 | Neutrophil |  |  |  |  |
| PTX3 | Neutrophil | detected in airway proteome | 20061.8 |  |  |
| RAB11B | Neutrophil |  |  |  |  |
| RAB26 | Neutrophil |  |  |  |  |
| RAB43 | Neutrophil |  |  |  |  |
| RAI14 | Neutrophil |  |  |  |  |
| RALB | Neutrophil | detected in airway proteome | 24583.8 |  |  |
| RASGRP3 | Neutrophil |  |  |  |  |
| RETN | Neutrophil | detected in airway proteome | 15772.1 |  |  |
| RGS19 | Neutrophil |  |  |  |  |
| RHBDF2 | Neutrophil |  |  |  |  |
| RICTOR | Neutrophil |  |  |  |  |
| RMND1 | Neutrophil |  |  |  |  |
| ROCK1 | Neutrophil |  |  |  |  |
| RPA3OS | Neutrophil |  |  |  |  |
| RPAP2 | Neutrophil |  |  |  |  |
| RRAGA | Neutrophil |  |  |  |  |
| RTN3 | Neutrophil |  |  |  |  |
| RYR1 | Neutrophil |  |  |  |  |
| S100A8 | Neutrophil | detected in airway proteome | 803277.1 |  |  |
| SCYL3 | Neutrophil |  |  |  |  |
| SFR1 | Neutrophil |  |  |  |  |
| SIRPB1 | Neutrophil |  |  |  |  |
| SLC16A3 | Neutrophil |  |  |  |  |
| SLC2A3 | Neutrophil |  |  |  |  |
| SMEK2 | Neutrophil |  |  |  |  |
| SPCS3 | Neutrophil |  |  |  |  |
| SPTAN1 | Neutrophil | detected in airway proteome | 23674.4 |  |  |
| STBD1 | Neutrophil |  |  |  |  |
| STIM1 | Neutrophil | detected in airway proteome | 52266.1 |  |  |
| SYF2 | Neutrophil |  |  |  |  |
| TALDO1 | Neutrophil | detected in airway proteome | 180771.9 |  |  |
| TCEAL3 | Neutrophil |  |  |  |  |
| TCEANC2 | Neutrophil |  |  |  |  |
| THOC1 | Neutrophil |  |  |  |  |
| TIMP2 | Neutrophil |  |  |  |  |
| TM9SF2 | Neutrophil |  |  |  |  |
| TMOD2 | Neutrophil |  |  |  |  |
| TMX1 | Neutrophil | detected in airway proteome | 23576.7 |  |  |
| TNFRSF10C | Neutrophil |  |  |  |  |
| TSC22D4 | Neutrophil |  |  |  |  |
| TSPAN14 | Neutrophil |  |  |  |  |
| TTC39B | Neutrophil |  |  |  |  |
| TTK | Neutrophil |  |  |  |  |
| TUBGCP2 | Neutrophil |  |  |  |  |
| UBL3 | Neutrophil |  |  |  |  |
| UQCRB | Neutrophil | detected in airway proteome | 116798.3 |  |  |
| VAMP8 | Neutrophil |  |  |  |  |
| VAT1 | Neutrophil | detected in airway proteome | 94620.8 |  |  |
| VKORC1L1 | Neutrophil |  |  |  |  |
| VNN2 | Neutrophil | detected in airway proteome | 16628.2 |  |  |
| VPS37B | Neutrophil | detected in airway proteome | 8694.7 |  |  |
| VPS72 | Neutrophil |  |  |  |  |
| WAS | Neutrophil |  |  |  |  |
| WBSCR16 | Neutrophil |  |  |  |  |
| WIPF1 | Neutrophil |  |  |  |  |
| YTHDF3 | Neutrophil |  |  |  |  |
| ZDHHC18 | Neutrophil |  |  |  |  |
| ZFP36L2 | Neutrophil |  |  |  |  |
| ZFYVE28 | Neutrophil |  |  |  |  |
| ZMYM3 | Neutrophil |  |  |  |  |
| ZNF106 | Neutrophil |  |  |  |  |
| ZNF281 | Neutrophil |  |  |  |  |
| ABCB1 | NK. |  |  |  |  |
| ABHD17A | NK. |  |  |  |  |
| AGK | NK. |  |  |  |  |
| AKAP5 | NK. |  |  |  |  |
| AKR1C3 | NK. | detected in airway proteome | 21379.4 |  |  |
| AOAH | NK. |  |  |  |  |
| APOBEC3G | NK. |  |  |  |  |
| B4GALT4 | NK. |  |  |  |  |
| CAPN5 | NK. |  |  |  |  |
| CD244 | NK. |  |  |  |  |
| CD247 | NK. |  |  |  |  |
| CD48 | NK. |  |  |  |  |
| CTSW | NK. |  |  |  |  |
| DHCR24 | NK. |  |  |  |  |
| DPF3 | NK. |  |  |  |  |
| ENPP1 | NK. |  |  |  |  |
| EOMES | NK. |  |  |  |  |
| ERMP1 | NK. |  |  |  |  |
| FCGR3A | NK. |  |  |  |  |
| FCGR3B | NK. | detected in airway proteome | 9161.7 |  |  |
| GNAI1 | NK. |  |  |  |  |
| GNLY | NK. |  |  |  |  |
| GOLGA8R | NK. |  |  |  |  |
| GOLIM4 | NK. |  |  |  |  |
| GOPC | NK. |  |  |  |  |
| GPR114 | NK. |  |  |  |  |
| GZMA | NK. |  |  |  |  |
| GZMB | NK. |  |  |  |  |
| GZMH | NK. |  |  |  |  |
| HAVCR2 | NK. |  |  |  |  |
| HDGFRP3 | NK. |  |  |  |  |
| IGFBP7 | NK. |  |  |  |  |
| IL18R1 | NK. |  |  |  |  |
| IL2RB | NK. |  |  |  |  |
| IRF2BPL | NK. |  |  |  |  |
| ITGA1 | NK. |  |  |  |  |
| KDELC1 | NK. |  |  |  |  |
| KIR2DS2 | NK. |  |  |  |  |
| KIR3DL1 | NK. |  |  |  |  |
| KIR3DL2 | NK. |  |  |  |  |
| KLRC1 | NK. |  |  |  |  |
| KLRD1 | NK. |  |  |  |  |
| LAT2 | NK. |  |  |  |  |
| LRG1 | NK. |  |  |  |  |
| MAFF | NK. |  |  |  |  |
| MATK | NK. |  |  |  |  |
| MCTP2 | NK. |  |  |  |  |
| NCALD | NK. |  |  |  |  |
| NCAM1 | NK. |  |  |  |  |
| NCR1 | NK. |  |  |  |  |
| NCR3 | NK. |  |  |  |  |
| NEIL1 | NK. |  |  |  |  |
| NFATC2 | NK. |  |  |  |  |
| ORAI2 | NK. |  |  |  |  |
| PNOC | NK. |  |  |  |  |
| PON2 | NK. | detected in airway proteome | 4735.4 |  |  |
| PRF1 | NK. |  |  |  |  |
| RGS3 | NK. |  |  |  |  |
| RUNX3 | NK. |  |  |  |  |
| SAMD3 | NK. |  |  |  |  |
| SH2D1B | NK. |  |  |  |  |
| SIGLEC7 | NK. |  |  |  |  |
| TBX21 | NK. |  |  |  |  |
| TNFRSF18 | NK. |  |  |  |  |
| TPST2 | NK. |  |  |  |  |
| TXK | NK. |  |  |  |  |
| AACS | NK.bright |  |  |  |  |
| ABCB1 | NK.bright |  |  |  |  |
| ACSL6 | NK.bright |  |  |  |  |
| AGK | NK.bright |  |  |  |  |
| ATHL1 | NK.bright |  |  |  |  |
| CD300A | NK.bright |  |  |  |  |
| CDH17 | NK.bright |  |  |  |  |
| CTSW | NK.bright |  |  |  |  |
| DENND1B | NK.bright |  |  |  |  |
| DPF3 | NK.bright |  |  |  |  |
| ENPP1 | NK.bright |  |  |  |  |
| ERMP1 | NK.bright |  |  |  |  |
| FUT7 | NK.bright |  |  |  |  |
| GNAI1 | NK.bright |  |  |  |  |
| GNLY | NK.bright |  |  |  |  |
| GOLIM4 | NK.bright |  |  |  |  |
| HDGFRP3 | NK.bright |  |  |  |  |
| IL18R1 | NK.bright |  |  |  |  |
| IL2RB | NK.bright |  |  |  |  |
| IRF2BPL | NK.bright |  |  |  |  |
| ITGA1 | NK.bright |  |  |  |  |
| KLRC1 | NK.bright |  |  |  |  |
| KLRD1 | NK.bright |  |  |  |  |
| MAFF | NK.bright |  |  |  |  |
| MATK | NK.bright |  |  |  |  |
| MLC1 | NK.bright |  |  |  |  |
| NCAM1 | NK.bright |  |  |  |  |
| NCR1 | NK.bright |  |  |  |  |
| NCR3 | NK.bright |  |  |  |  |
| NEIL1 | NK.bright |  |  |  |  |
| PSD4 | NK.bright |  |  |  |  |
| RAB38 | NK.bright |  |  |  |  |
| RGS3 | NK.bright |  |  |  |  |
| RUNX3 | NK.bright |  |  |  |  |
| SARDH | NK.bright |  |  |  |  |
| STARD10 | NK.bright |  |  |  |  |
| TNFRSF18 | NK.bright |  |  |  |  |
| TPST2 | NK.bright |  |  |  |  |
| XCL2 | NK.bright |  |  |  |  |
| ABHD17A | NK.dim |  |  |  |  |
| AKR1C3 | NK.dim | detected in airway proteome | 21379.4 |  |  |
| B4GALT4 | NK.dim |  |  |  |  |
| CAPN5 | NK.dim |  |  |  |  |
| CD160 | NK.dim |  |  |  |  |
| CD247 | NK.dim |  |  |  |  |
| CD48 | NK.dim |  |  |  |  |
| CDKN2A | NK.dim |  |  |  |  |
| CST7 | NK.dim |  |  |  |  |
| F2R | NK.dim |  |  |  |  |
| FCGR3A | NK.dim |  |  |  |  |
| FGFBP2 | NK.dim |  |  |  |  |
| FUT8 | NK.dim |  |  |  |  |
| GNLY | NK.dim |  |  |  |  |
| GOLGA8R | NK.dim |  |  |  |  |
| IDS | NK.dim |  |  |  |  |
| IGF2R | NK.dim |  |  |  |  |
| IL2RB | NK.dim |  |  |  |  |
| KIAA1671 | NK.dim |  |  |  |  |
| KIR2DL1 | NK.dim |  |  |  |  |
| KIR2DS2 | NK.dim |  |  |  |  |
| KIR3DL1 | NK.dim |  |  |  |  |
| KIR3DL2 | NK.dim |  |  |  |  |
| KPNA5 | NK.dim |  |  |  |  |
| NCAM1 | NK.dim |  |  |  |  |
| NFATC2 | NK.dim |  |  |  |  |
| PLEKHF1 | NK.dim |  |  |  |  |
| PNOC | NK.dim |  |  |  |  |
| PRF1 | NK.dim |  |  |  |  |
| RGS3 | NK.dim |  |  |  |  |
| SH2D1B | NK.dim |  |  |  |  |
| SYTL2 | NK.dim |  |  |  |  |
| TBX21 | NK.dim |  |  |  |  |
| C9orf78 | nTregs |  |  |  |  |
| CD3D | nTregs |  |  |  |  |
| CD5 | nTregs |  |  |  |  |
| CPN2 | nTregs |  |  |  |  |
| CXCR4 | nTregs |  |  |  |  |
| EVPL | nTregs | detected in airway proteome | 370133.6 |  |  |
| FOXP3 | nTregs |  |  |  |  |
| GPA33 | nTregs |  |  |  |  |
| KLKB1 | nTregs |  |  |  |  |
| MLLT3 | nTregs |  |  |  |  |
| NECAP2 | nTregs |  |  |  |  |
| PGM2L1 | nTregs |  |  |  |  |
| PRPSAP2 | nTregs | detected in airway proteome | 24365.2 |  |  |
| RAB28 | nTregs |  |  |  |  |
| SIT1 | nTregs |  |  |  |  |
| WHSC1L1 | nTregs |  |  |  |  |
| ACY3 | pDC |  |  |  |  |
| ADA | pDC |  |  |  |  |
| ALG2 | pDC |  |  |  |  |
| B4GALT5 | pDC |  |  |  |  |
| C12orf45 | pDC |  |  |  |  |
| CCDC50 | pDC |  |  |  |  |
| CD2AP | pDC |  |  |  |  |
| CDH1 | pDC |  |  |  |  |
| CLIC3 | pDC | detected in airway proteome | 188172.7 |  |  |
| CRYM | pDC |  |  |  |  |
| CXorf21 | pDC |  |  |  |  |
| DDAH2 | pDC |  |  |  |  |
| DGKZ | pDC |  |  |  |  |
| DHTKD1 | pDC |  |  |  |  |
| DNAJB4 | pDC |  |  |  |  |
| ENDOG | pDC | detected in airway proteome | 16148.0 |  |  |
| EPHA2 | pDC |  |  |  |  |
| ERN1 | pDC |  |  |  |  |
| FAM213A | pDC | detected in airway proteome | 10756.5 |  |  |
| FCHSD2 | pDC |  |  |  |  |
| FLNB | pDC | detected in airway proteome | 310540.1 |  |  |
| HYOU1 | pDC | detected in airway proteome | 160839.1 |  |  |
| IDH3A | pDC | detected in airway proteome | 63826.7 |  |  |
| IDH3G | pDC | detected in airway proteome | 22190.9 |  |  |
| IL3RA | pDC |  |  |  |  |
| IRF7 | pDC |  |  |  |  |
| IRF8 | pDC |  |  |  |  |
| KRTCAP2 | pDC |  |  |  |  |
| LGMN | pDC |  |  |  |  |
| MAP1A | pDC |  |  |  |  |
| MAP2K6 | pDC |  |  |  |  |
| MCC | pDC |  |  |  |  |
| NGLY1 | pDC |  |  |  |  |
| NPC1 | pDC |  |  |  |  |
| PACSIN1 | pDC |  |  |  |  |
| PALD1 | pDC |  |  |  |  |
| PARP10 | pDC |  |  |  |  |
| PFKFB2 | pDC |  |  |  |  |
| PIK3R5 | pDC |  |  |  |  |
| PLEK | pDC | detected in airway proteome | 6435.7 |  |  |
| PPP1R14B | pDC |  |  |  |  |
| PRKRA | pDC |  |  |  |  |
| PTGDS | pDC |  |  |  |  |
| PTPRS | pDC |  |  |  |  |
| RAB2B | pDC |  |  |  |  |
| RRBP1 | pDC | detected in airway proteome | 14532.1 |  |  |
| SCAMP5 | pDC |  |  |  |  |
| SMIM5 | pDC |  |  |  |  |
| SPCS2 | pDC | detected in airway proteome | 14394.7 |  |  |
| SRGAP2 | pDC |  |  |  |  |
| TBC1D4 | pDC |  |  |  |  |
| TLR7 | pDC |  |  |  |  |
| TRAF4 | pDC |  |  |  |  |
| TXNDC5 | pDC | detected in airway proteome | 103227.6 |  |  |
| XIRP1 | pDC |  |  |  |  |
| ZFYVE26 | pDC |  |  |  |  |
| ZNRF2 | pDC |  |  |  |  |
| ANK3 | T4 |  |  |  |  |
| APOA4 | T4 | detected in airway proteome | 11921.2 |  |  |
| BAG3 | T4 | detected in airway proteome | 34498.1 |  |  |
| BCL11B | T4 |  |  |  |  |
| CCNT1 | T4 |  |  |  |  |
| CD109 | T4 |  |  |  |  |
| CD3E | T4 |  |  |  |  |
| CD3G | T4 |  |  |  |  |
| CD4 | T4 |  |  |  |  |
| CD5 | T4 |  |  |  |  |
| CDK1 | T4 |  |  |  |  |
| CKAP5 | T4 |  |  |  |  |
| CLK4 | T4 |  |  |  |  |
| DDX21 | T4 |  |  |  |  |
| EIF3C | T4 |  |  |  |  |
| EPHX2 | T4 |  |  |  |  |
| FKBP8 | T4 | detected in airway proteome | 6738.2 |  |  |
| GINS3 | T4 |  |  |  |  |
| KEAP1 | T4 |  |  |  |  |
| KIF15 | T4 |  |  |  |  |
| LIN37 | T4 |  |  |  |  |
| MBD1 | T4 |  |  |  |  |
| MINA | T4 |  |  |  |  |
| MRTO4 | T4 |  |  |  |  |
| NDRG3 | T4 |  |  |  |  |
| NDST1 | T4 |  |  |  |  |
| NOP14 | T4 |  |  |  |  |
| NOSIP | T4 |  |  |  |  |
| PBK | T4 |  |  |  |  |
| PRDX2 | T4 | detected in airway proteome | 91991.6 |  |  |
| PRMT3 | T4 |  |  |  |  |
| PSAT1 | T4 |  |  |  |  |
| PSMG1 | T4 |  |  |  |  |
| RBL2 | T4 |  |  |  |  |
| RGCC | T4 |  |  |  |  |
| RLTPR | T4 |  |  |  |  |
| RRP1B | T4 |  |  |  |  |
| SPAG5 | T4 |  |  |  |  |
| SQSTM1 | T4 |  |  |  |  |
| STAU1 | T4 |  |  |  |  |
| TCRBV5S1A1T | T4 |  |  |  |  |
| THEMIS | T4 |  |  |  |  |
| TNFRSF4 | T4 |  |  |  |  |
| TRADD | T4 | detected in airway proteome | 9496.1 |  |  |
| TRAT1 | T4 |  |  |  |  |
| UHRF1BP1 | T4 |  |  |  |  |
| UTP3 | T4 |  |  |  |  |
| WHSC1 | T4 |  |  |  |  |
| ZBTB7B | T4 |  |  |  |  |
| FAM26E | T4.CM |  |  |  |  |
| FASTKD5 | T4.CM |  |  |  |  |
| FKBP8 | T4.CM | detected in airway proteome | 6738.2 |  |  |
| KNTC1 | T4.CM |  |  |  |  |
| PUM1 | T4.CM |  |  |  |  |
| RANGRF | T4.CM |  |  |  |  |
| ZNF688 | T4.CM |  |  |  |  |
| ABCF2 | T4.EM |  |  |  |  |
| CD40LG | T4.EM |  |  |  |  |
| CETN3 | T4.EM |  |  |  |  |
| CLUH | T4.EM |  |  |  |  |
| CNDP1 | T4.EM |  |  |  |  |
| DDX21 | T4.EM |  |  |  |  |
| DNAJC7 | T4.EM |  |  |  |  |
| FETUB | T4.EM |  |  |  |  |
| HDLBP | T4.EM | detected in airway proteome | 15257.7 |  |  |
| HSP90AB1 | T4.EM | detected in airway proteome | 165953.6 |  |  |
| IPO11 | T4.EM |  |  |  |  |
| KIAA0020 | T4.EM |  |  |  |  |
| NOL7 | T4.EM |  |  |  |  |
| NOP14 | T4.EM |  |  |  |  |
| PDCD11 | T4.EM |  |  |  |  |
| PSMG1 | T4.EM |  |  |  |  |
| RANBP1 | T4.EM | detected in airway proteome | 41574.0 |  |  |
| RNF146 | T4.EM |  |  |  |  |
| RPL12 | T4.EM | detected in airway proteome | 103997.4 |  |  |
| RPL17 | T4.EM | detected in airway proteome | 36790.4 |  |  |
| RPL27 | T4.EM | detected in airway proteome | 115404.9 |  |  |
| RPL38 | T4.EM | detected in airway proteome | 11092.7 |  |  |
| RSL1D1 | T4.EM |  |  |  |  |
| SERPINA10 | T4.EM |  |  |  |  |
| SHBG | T4.EM |  |  |  |  |
| SPAG5 | T4.EM |  |  |  |  |
| WDR43 | T4.EM |  |  |  |  |
| YBX1 | T4.EM | detected in airway proteome | 2624.0 |  |  |
| APOA4 | T4.EMRA | detected in airway proteome | 11921.2 |  |  |
| C1S | T4.EMRA |  |  |  |  |
| CCDC85B | T4.EMRA |  |  |  |  |
| CFH | T4.EMRA |  |  |  |  |
| CFHR5 | T4.EMRA |  |  |  |  |
| CNDP1 | T4.EMRA |  |  |  |  |
| DDX21 | T4.EMRA |  |  |  |  |
| EIF5A | T4.EMRA | detected in airway proteome | 70184.0 |  |  |
| FETUB | T4.EMRA |  |  |  |  |
| GHITM | T4.EMRA |  |  |  |  |
| GPLD1 | T4.EMRA |  |  |  |  |
| GRPEL1 | T4.EMRA | detected in airway proteome | 12184.4 |  |  |
| HSP90AB4P | T4.EMRA |  |  |  |  |
| KIF2C | T4.EMRA |  |  |  |  |
| PCDH11X | T4.EMRA |  |  |  |  |
| PNPO | T4.EMRA |  |  |  |  |
| PSMG1 | T4.EMRA |  |  |  |  |
| RANBP1 | T4.EMRA | detected in airway proteome | 41574.0 |  |  |
| RNF213 | T4.EMRA | detected in airway proteome | 25926.5 |  |  |
| RPL12 | T4.EMRA | detected in airway proteome | 103997.4 |  |  |
| SERPINA10 | T4.EMRA |  |  |  |  |
| SERPINA5 | T4.EMRA |  |  |  |  |
| SHBG | T4.EMRA |  |  |  |  |
| TTR | T4.EMRA | detected in airway proteome | 27730.9 |  |  |
| YBX1 | T4.EMRA | detected in airway proteome | 2624.0 |  |  |
| AVEN | T4.naive |  |  |  |  |
| BCL11B | T4.naive |  |  |  |  |
| CLK4 | T4.naive |  |  |  |  |
| DENND2D | T4.naive |  |  |  |  |
| EPHX2 | T4.naive |  |  |  |  |
| POTEE | T4.naive |  |  |  |  |
| PSIP1 | T4.naive |  |  |  |  |
| TCRBV5S1A1T | T4.naive |  |  |  |  |
| TMEM254 | T4.naive |  |  |  |  |
| AATF | T8 |  |  |  |  |
| ABCB1 | T8 |  |  |  |  |
| ACN9 | T8 |  |  |  |  |
| ADAT1 | T8 |  |  |  |  |
| ATHL1 | T8 |  |  |  |  |
| BCL11B | T8 |  |  |  |  |
| BTN3A3 | T8 |  |  |  |  |
| CBLB | T8 |  |  |  |  |
| CCL5 | T8 |  |  |  |  |
| CD3D | T8 |  |  |  |  |
| CD3E | T8 |  |  |  |  |
| CD3EAP | T8 |  |  |  |  |
| CD3G | T8 |  |  |  |  |
| CD5 | T8 |  |  |  |  |
| CD6 | T8 |  |  |  |  |
| CD8A | T8 |  |  |  |  |
| CD8B | T8 |  |  |  |  |
| CLIP4 | T8 |  |  |  |  |
| CST7 | T8 |  |  |  |  |
| CTSW | T8 |  |  |  |  |
| DBN1 | T8 |  |  |  |  |
| DDX24 | T8 |  |  |  |  |
| DHX33 | T8 |  |  |  |  |
| EOMES | T8 |  |  |  |  |
| FKBP11 | T8 | detected in airway proteome | 40465.3 |  |  |
| GLS | T8 |  |  |  |  |
| GPR171 | T8 |  |  |  |  |
| GRAP2 | T8 |  |  |  |  |
| GZMA | T8 |  |  |  |  |
| GZMB | T8 |  |  |  |  |
| GZMH | T8 |  |  |  |  |
| GZMK | T8 |  |  |  |  |
| GZMM | T8 |  |  |  |  |
| HIST1H3A | T8 | detected in airway proteome | 3714.0 |  |  |
| IGSF8 | T8 |  |  |  |  |
| IL2RB | T8 |  |  |  |  |
| KLRG1 | T8 |  |  |  |  |
| KLRK1 | T8 |  |  |  |  |
| LCK | T8 |  |  |  |  |
| LETMD1 | T8 |  |  |  |  |
| LIMA1 | T8 |  |  |  |  |
| LIPT2 | T8 |  |  |  |  |
| LYAR | T8 |  |  |  |  |
| MECP2 | T8 |  |  |  |  |
| MLLT11 | T8 |  |  |  |  |
| MLLT6 | T8 |  |  |  |  |
| NCALD | T8 |  |  |  |  |
| NIPSNAP1 | T8 | detected in airway proteome | 17503.0 |  |  |
| NT5E | T8 |  |  |  |  |
| PRF1 | T8 |  |  |  |  |
| PTPRCAP | T8 |  |  |  |  |
| PTRH1 | T8 |  |  |  |  |
| SCCPDH | T8 | detected in airway proteome | 18099.7 |  |  |
| SEMA4B | T8 |  |  |  |  |
| SH2D2A | T8 |  |  |  |  |
| SLC25A4 | T8 |  |  |  |  |
| SLC39A14 | T8 |  |  |  |  |
| SUPV3L1 | T8 |  |  |  |  |
| TBX21 | T8 |  |  |  |  |
| THEMIS | T8 |  |  |  |  |
| TRAT1 | T8 |  |  |  |  |
| UBASH3A | T8 |  |  |  |  |
| VWA8 | T8 |  |  |  |  |
| YBX3 | T8 |  |  |  |  |
| ZBTB44 | T8 |  |  |  |  |
| CD226 | T8.CM |  |  |  |  |
| CD8A | T8.CM |  |  |  |  |
| CD8B | T8.CM |  |  |  |  |
| LAX1 | T8.CM |  |  |  |  |
| SLC29A1 | T8.CM |  |  |  |  |
| SLC43A3 | T8.CM |  |  |  |  |
| TRBV19 | T8.CM |  |  |  |  |
| BTN3A3 | T8.EM |  |  |  |  |
| C9orf142 | T8.EM |  |  |  |  |
| CCL5 | T8.EM |  |  |  |  |
| CD8A | T8.EM |  |  |  |  |
| CD8B | T8.EM |  |  |  |  |
| CLIP4 | T8.EM |  |  |  |  |
| CST7 | T8.EM |  |  |  |  |
| CTSW | T8.EM |  |  |  |  |
| DBN1 | T8.EM |  |  |  |  |
| ENPP4 | T8.EM |  |  |  |  |
| EOMES | T8.EM |  |  |  |  |
| GNLY | T8.EM |  |  |  |  |
| GPR171 | T8.EM |  |  |  |  |
| GPR56 | T8.EM |  |  |  |  |
| GZMA | T8.EM |  |  |  |  |
| GZMH | T8.EM |  |  |  |  |
| GZMK | T8.EM |  |  |  |  |
| GZMM | T8.EM |  |  |  |  |
| IKZF3 | T8.EM |  |  |  |  |
| KIF21A | T8.EM |  |  |  |  |
| KLRG1 | T8.EM |  |  |  |  |
| MCTP2 | T8.EM |  |  |  |  |
| MLLT6 | T8.EM |  |  |  |  |
| MXRA7 | T8.EM |  |  |  |  |
| PRF1 | T8.EM |  |  |  |  |
| PTPRCAP | T8.EM |  |  |  |  |
| PYHIN1 | T8.EM |  |  |  |  |
| RABGAP1L | T8.EM |  |  |  |  |
| S1PR5 | T8.EM |  |  |  |  |
| SLC25A4 | T8.EM |  |  |  |  |
| SYNE2 | T8.EM |  |  |  |  |
| SYTL2 | T8.EM |  |  |  |  |
| TNS1 | T8.EM |  |  |  |  |
| TRGC1 | T8.EM |  |  |  |  |
| ZBTB38 | T8.EM |  |  |  |  |
| CD8A | T8.EMRA |  |  |  |  |
| CRTAM | T8.EMRA |  |  |  |  |
| DBN1 | T8.EMRA |  |  |  |  |
| EOMES | T8.EMRA |  |  |  |  |
| GZMH | T8.EMRA |  |  |  |  |
| HOOK1 | T8.EMRA |  |  |  |  |
| IDS | T8.EMRA |  |  |  |  |
| KIF21A | T8.EMRA |  |  |  |  |
| NAPB | T8.EMRA |  |  |  |  |
| PDLIM2 | T8.EMRA | detected in airway proteome | 22565.4 |  |  |
| PIP4K2A | T8.EMRA |  |  |  |  |
| PLEKHA1 | T8.EMRA |  |  |  |  |
| PTPRCAP | T8.EMRA |  |  |  |  |
| RASSF1 | T8.EMRA |  |  |  |  |
| S1PR5 | T8.EMRA |  |  |  |  |
| TMEM143 | T8.EMRA |  |  |  |  |
| TNFSF4 | T8.EMRA |  |  |  |  |
| TNS1 | T8.EMRA |  |  |  |  |
| UBLCP1 | T8.EMRA |  |  |  |  |
| ADCK3 | T8.naive |  |  |  |  |
| ATM | T8.naive |  |  |  |  |
| BCL2 | T8.naive |  |  |  |  |
| CD27 | T8.naive |  |  |  |  |
| CD8A | T8.naive |  |  |  |  |
| CD8B | T8.naive |  |  |  |  |
| CENPV | T8.naive |  |  |  |  |
| EIF4EBP3 | T8.naive |  |  |  |  |
| GNG8 | T8.naive |  |  |  |  |
| HIC1 | T8.naive |  |  |  |  |
| LEF1 | T8.naive |  |  |  |  |
| LMO7 | T8.naive | detected in airway proteome | 112273.3 |  |  |
| NAA16 | T8.naive |  |  |  |  |
| NR3C2 | T8.naive |  |  |  |  |
| NT5E | T8.naive |  |  |  |  |
| PDK1 | T8.naive |  |  |  |  |
| PNMA3 | T8.naive |  |  |  |  |
| PPAP2A | T8.naive |  |  |  |  |
| SELH | T8.naive |  |  |  |  |
| SH3BGRL2 | T8.naive | detected in airway proteome | 27152.1 |  |  |
| THEMIS | T8.naive |  |  |  |  |
| THNSL1 | T8.naive |  |  |  |  |
| THYN1 | T8.naive |  |  |  |  |
| WDR89 | T8.naive |  |  |  |  |
| ANK3 | Th1 |  |  |  |  |
| CD6 | Th1 |  |  |  |  |
| MPRIP | Th1 |  |  |  |  |
| C16orf74 | Th17 |  |  |  |  |
| CA5B | Th17 |  |  |  |  |
| MPRIP | Th17 |  |  |  |  |
| N4BP3 | Th17 |  |  |  |  |
| SUFU | Th17 |  |  |  |  |
| TPPP | Th17 |  |  |  |  |
| COQ10A | Th2 |  |  |  |  |
| SARAF | Th2 |  |  |  |  |
| SKAP1 | Th2 |  |  |  |  |
| AQR | Treg |  |  |  |  |
| C9orf78 | Treg |  |  |  |  |
| CCM2 | Treg |  |  |  |  |
| CD27 | Treg |  |  |  |  |
| CD3D | Treg |  |  |  |  |
| CD3E | Treg |  |  |  |  |
| CD4 | Treg |  |  |  |  |
| CD5 | Treg |  |  |  |  |
| CEACAM1 | Treg | detected in airway proteome | 19325.9 |  |  |
| CFI | Treg |  |  |  |  |
| CPN2 | Treg |  |  |  |  |
| CXCR4 | Treg |  |  |  |  |
| EFEMP1 | Treg |  |  |  |  |
| EPB41 | Treg |  |  |  |  |
| EVPL | Treg | detected in airway proteome | 370133.6 |  |  |
| FAM120B | Treg |  |  |  |  |
| FOXO1 | Treg |  |  |  |  |
| FOXP3 | Treg |  |  |  |  |
| GCC2 | Treg |  |  |  |  |
| GPA33 | Treg |  |  |  |  |
| GPX3 | Treg |  |  |  |  |
| IKZF2 | Treg |  |  |  |  |
| IKZF4 | Treg |  |  |  |  |
| ITGA6 | Treg |  |  |  |  |
| ITIH4 | Treg | detected in airway proteome | 23811.3 |  |  |
| KLKB1 | Treg |  |  |  |  |
| LAX1 | Treg |  |  |  |  |
| MCRS1 | Treg |  |  |  |  |
| MEOX1 | Treg |  |  |  |  |
| MLLT3 | Treg |  |  |  |  |
| NECAP2 | Treg |  |  |  |  |
| NIPBL | Treg | detected in airway proteome | 37022.4 |  |  |
| PGM2L1 | Treg |  |  |  |  |
| PI16 | Treg |  |  |  |  |
| PLCL1 | Treg |  |  |  |  |
| PNN | Treg |  |  |  |  |
| RBBP6 | Treg |  |  |  |  |
| SIT1 | Treg |  |  |  |  |
| SLC35D1 | Treg |  |  |  |  |
| SMC1A | Treg |  |  |  |  |
| SMC3 | Treg |  |  |  |  |
| SPOCK2 | Treg |  |  |  |  |
| TBC1D4 | Treg |  |  |  |  |
| TRIM56 | Treg |  |  |  |  |
| WDR33 | Treg |  |  |  |  |
| WHSC1L1 | Treg |  |  |  |  |
| ZC3H12D | Treg |  |  |  |  |
